# Interleukin-34 and debris clearance by mononuclear phagocytes drive retinal pigment epithelium regeneration in zebrafish

**DOI:** 10.1101/2025.01.10.632236

**Authors:** Lyndsay L. Leach, Rebecca G. Gonzalez, Sayuri U. Jayawardena, Jeffrey M. Gross

## Abstract

The retinal pigment epithelium (RPE) surrounds the posterior eye and maintains the health and function of the photoreceptors. Consequently, RPE dysfunction or damage has a devastating impact on vision. Due to complex etiologies, there are currently no cures for patients with RPE degenerative diseases, which remain some of the most prevalent causes of vision loss worldwide. Further, owing to a limited capacity for mammalian tissue repair, we know little about how the RPE regenerates. Here, we utilize zebrafish as a model to uncover novel mechanisms driving intrinsic RPE regeneration. We show that interleukin-34 signaling from damaged RPE is required for precisely timed recruitment of mononuclear phagocytes (MNPs) to the injury site. Additionally, we find that cellular debris clearance by MNPs is indispensable for regeneration, as microglia-deficient zebrafish fail to regenerate RPE and photoreceptor tissues. Together, our results establish specific pro-regenerative functions of MNPs after RPE damage.

## Introduction

The retinal pigment epithelium (RPE) is a multifunctional monolayer located in the posterior eye at the interface between the choroid vasculature and the light-sensing photoreceptors of the retina. The RPE comprises the blood-retinal-barrier and forms a tightly interdigitated functional unit with the photoreceptors, which rely on the RPE for nutrients, growth factors, and maintenance of excitability, among other things^1^. Unsurprisingly, damage to the RPE has devastating consequences for photoreceptor function and for vision in general. Examples include RPE degenerative diseases, such as age-related macular degeneration (AMD) and Stargardt disease, where primary insult to the RPE yields secondary photoreceptor degeneration and eventual blindness. Indeed, RPE degenerative diseases remain some of the most prevalent causes of vision loss worldwide, and AMD alone is estimated to affect 288 million people by 2040^2^. Owing to the limited reparative capacity of mature mammalian RPE^3^, there is a major knowledge gap linking RPE damage and the mechanisms underlying intrinsic repair and regeneration. We developed and characterized a zebrafish RPE ablation paradigm^4^, which, due to the robust regenerative capacity of this vertebrate model, has allowed us to elucidate novel mechanisms underlying the intrinsic RPE regenerative response.

Fitting a common theme among zebrafish tissue regeneration studies^5–7^, we found that elements of the innate immune response play a central, pro-regenerative role after RPE damage^8–10^. Specifically, we demonstrated temporal injury site infiltration dynamics and general functional requirements for mononuclear phagocytes (MNPs; i.e., microglia, monocyte-derived or tissue macrophages) during RPE regeneration; and bulk RNA-sequencing (RNA-seq) experiments were instrumental in uncovering immune-related candidate genes and pathways that may regulate RPE regeneration^8–10^, many of which remain to be investigated. One interesting candidate is the cytokine, interleukin-34 (IL-34), which is expressed by the RPE at early and peak regenerative timepoints^8^. IL-34 signals through colony stimulating factor 1 receptor (CSF1R; one of four known receptors^11–14^) to evoke a plethora of critical functions in MNPs responding to tissue damage or disease, including recruitment, proliferation, phagocytosis, and survival^15–17^. While the inflammatory or anti-inflammatory outcome of IL-34 signaling appears to be context specific, it has been shown to play a protective role in tissues of the central nervous system (CNS). For example, in the mouse retina, a distinct population of IL-34-dependent microglia (resident CNS macrophages) is recruited to the apical surface of the RPE and provides protection after acute photoreceptor damage^18^. In the zebrafish retina, we showed that pharmacologically inhibiting CSF1R impairs MNP function and RPE regeneration^8,9^. Additionally, in the brain, injection of recombinant IL-34 increased microglia densities, lowered soluble amyloid-ß peptide levels, and rescued cognitive function in a mouse model of Alzheimer’s disease (AD)^19^. Autocrine IL-34 signaling in neurons may also play a protective role against tissue degeneration in mouse AD-like brains^20^, highlighting the versatility and importance of this pathway in CNS tissues.

Despite the devastating impact of RPE degenerative diseases and in contrast to the many discoveries made in recent decades in the field of retinal regeneration^21,22^, we know comparatively little about intrinsic RPE regenerative mechanisms. Further, apart from the elegant work by O’Koren *et al.*^18^ to uncover the RPE-protective effects of IL-34-dependent microglia, we know almost nothing about the role of IL-34 signaling in RPE damage responses. Here, we explore the intrinsic regenerative capacity of the RPE in zebrafish with systemic loss of *il34*^23^ or *csf1ra/csf1rb*^24,25^, the latter of which is a microglia loss-of-function model. We find that *il34* is required for RPE regeneration and drives the precisely timed recruitment of MNPs to the RPE injury site. Further, we show that cellular debris clearance by MNPs is indispensable for regeneration, as mutants lacking microglia fail to regenerate RPE and photoreceptors in the long-term. Collectively, our results validate IL-34 as an RPE-derived pro-regenerative signal and spotlight the phagocytic role of MNPs as a critical determinant of RPE regeneration success.

## Results

### Pigment recovery is impaired in *il34* mutants during peak RPE regeneration

We previously detected upregulation of *il34* expression in regenerating RPE by RNA-seq^8^. *In situ* hybridization was utilized to validate these findings and confirm that *il34* was expressed specifically by the RPE (Figure S1A,B). Expression increased throughout the RPE and central injury site in ablated larvae at 2 days post-injury (dpi) (Figure S1B,D) compared to unablated controls (Figure S1A,C). To determine the consequence of IL-34 signaling loss on RPE regeneration, we crossed *il34* mutant zebrafish^23^ to our RPE genetic ablation line, *Tg(rpe65a:nfsB-eGFP),* and examined tissues at 4 dpi, which we previously characterized as a peak regenerative timepoint^4^. This transgenic line enables targeted ablation of mature RPE utilizing *rpe65a* enhancer-driven expression of nitroreductase *(nfsB)* fused to eGFP for visualization. Nitroreductase activates and converts metronidazole (MTZ) into a DNA-crosslinking agent to promote apoptosis in transgene-expressing cells, and the efficacy of this approach has been demonstrated in many tissue types^4, 26–29^. For all RPE ablation experiments in this study, larvae sustained a 24-hour immersion treatment with 10 mM MTZ from 5 – 6 days post-fertilization (dpf) and were subsequently allowed to recover post-injury until fixation. We quantified central RPE/pigment recovery using RpEGEN^30^, which revealed large regions of the *il34*^-/-^ RPE ablation site with significantly less overall pigment when compared to *il34*^+/+^ sibling controls (Figure 1A,B). This included two dorsal–central swathes spanning a total of ∼78-degrees, collectively representing ∼65% of the central RPE (Figure 1A,B; *light blue boxes*). Indeed, *il34*^-/-^ larval sections showed distinct central deficits in the RPE layer, which presented as a lack of organized pigmented tissue regrowth and, instead, retention of lightly pigmented clumps compared to *il34*^+/+^ siblings (Figure 1C,D). The significance of this finding is underscored by the apparent dispensability of *il34* for normal RPE development and maintenance, as unablated *il34^-/-^*RPE were phenotypically normal, and displayed defined apical microvilli and robust eGFP expression at 6 dpf and 9 dpf (Figure S2A–D). Importantly, TUNEL labeling revealed no significant differences in apoptotic signatures between unablated (6 dpf) or ablated (1 dpi) *il34*^+/+^ and *il34*^-/-^ siblings (Figure S3A–C), indicating loss of *il34* does not affect baseline cell survival or the severity of the initial ablation, respectively. Together, these findings show that *il34* expression is restricted to the RPE, and that systemic loss of *il34* impedes RPE repair at a peak regenerative timepoint; thus, establishing IL-34 signaling as a critical pathway for successful RPE regeneration.

**Figure 1.**
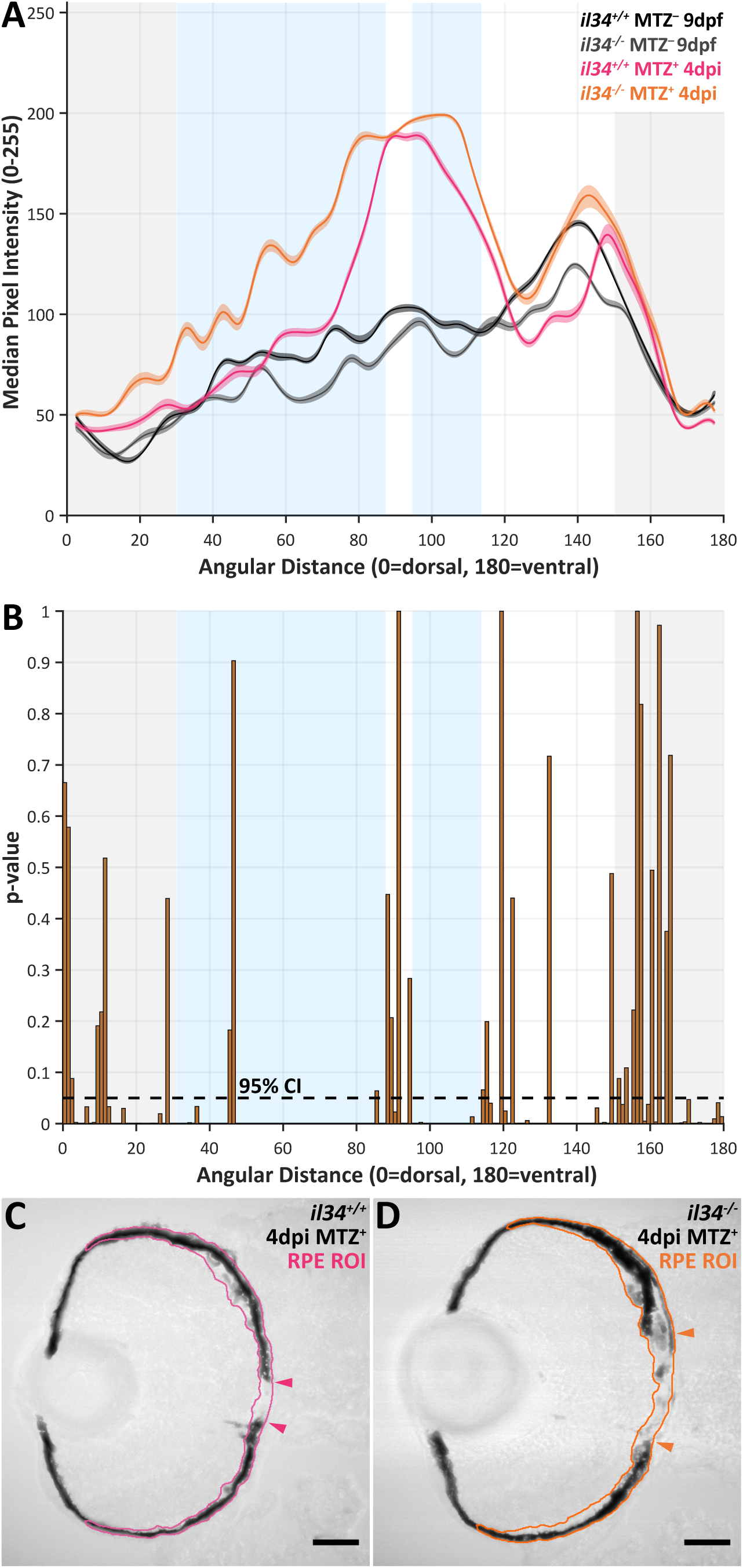
RPE regeneration is impaired in *il34* mutants at 4 dpi. (A,B) RpEGEN output plots showing **(A)** median pixel intensity values (8-bit grayscale) and **(B)** p-values spanning the dorsal (0°) to ventral (180°) angular distance of the RPE. *Black dotted line* represents p=0.05. **(A)** Ablated (MTZ^+^) *il34^-/-^* larvae *(orange line)* show higher (lighter/less pigmented) pixel intensities across the central two-thirds of the RPE compared to any other dataset. **(A,B)** *Light blue boxes* highlight large central regions where pigment recovery is significantly impaired. *Gray boxes* designate the peripheral third of the RPE, which remains intact in this ablation model. **(C,D)** Transverse confocal micrographs showing brightfield acquisitions of 4 dpi ablated (MTZ^+^) **(C)** *il34^+/+^* and **(D)** *il34^-/-^* larvae. *Orange and pink outlines* encompass RPE regions of interest (ROI). *Arrowheads* designate the central-most edges of intact RPE. Scale bars = 40µm. Experimental replicates and statistical information can be found in Table S1.

### Mononuclear phagocyte recruitment and debris clearance are delayed in *il34* mutants at early regeneration timepoints

In addition to identifying RPE-specific *il34* upregulation, we also previously showed that MNPs are robustly recruited to the RPE at 2 and 3 dpi and are functionally required for RPE regeneration^8^. It is known that the IL-34–CSF1R signaling axis promotes MNP recruitment in physiological^31^ and pathological^32^ contexts. To begin to establish a mechanism by which IL-34 signaling impacts RPE regeneration, we utilized a transgenic reporter line *(mpeg1:mCherry)*^33^ and an antibody to lgals3bpb (4C4)^34,35^ to visually assess MNP recruitment in *il34* mutants. Importantly, the specificity of these markers for distinguishing zebrafish tissue resident macrophages from monocyte-derived lineages has been unclear, although a recent study showed that lgals3bpb/4C4 expression is largely restricted to CNS microglia under homeostatic conditions^35^. For comprehensive purposes, however, we will refer to mCherry^+^ and 4C4^+^ cells as MNPs throughout the discussion of IL-34-related results. Prior characterization of *il34* mutants found systemic and retina-specific MNP tissue seeding deficiencies at embryonic and early larval stages (≤5 dpf)^23,31^, though systemic numbers eventually resolved to match wild type densities by 8 dpf^23^. Consistently, here, analysis of ≥7 dpf larvae showed resident microglia were present in unablated *il34^+/+^* and *il34^-/-^* larval eyes, and there were no significant differences in 4C4 or mCherry labeling between sibling groups (Figure S4A–F). In ablated animals, however, quantification of both 4C4 and mCherry showed significantly less MNPs in the RPE layer in *il34^-/-^* larvae at 2 dpi (Figure 2A–D). By 3 dpi, RPE-localized mCherry levels were similar between *il34^+/+^* and *il34^-/-^*siblings (Figure 2E–G), indicating that loss of *il34* causes a delay in MNP recruitment. Despite normalized MNP infiltration at 3 dpi, we observed marked retention of eGFP^+^ debris in *il34^-/-^* RPE tissues (Figure 2F), which was significantly increased when compared to wild type siblings (Figure 2H). Interestingly, statistical comparisons among timepoints within the *il34^+/+^* cohort revealed a significant drop in eGFP*^+^* debris from 2 – 3 dpi, and again from 3 – 4 dpi (Figure 2H; *solid gray significance bars*). Conversely, in *il34^-/-^* animals, eGFP signals remained similarly elevated between 2 – 3 dpi and then dropped significantly from 3 – 4 dpi (Figure 2H; *dotted gray significance bars*). These data suggest that 2 – 3 dpi is an important time window during which recruited MNPs clean up the injury site to prompt subsequent regenerative processes. In *il34^-/-^* larvae, delayed recruitment of MNPs likely results in a complementary shift in debris clearance to 3 – 4 dpi, as there was no significant difference between 4 dpi wild type and mutant sibling eGFP levels (Figure 2H). To evaluate active phagocytosis, we measured eGFP^+^ particle internalization by mCherry^+^ cells using Imaris. As an example, Figure S5B highlights an *il34^+/+^* MNP with superficial *(red)* and internalized *(blue* and *purple)* eGFP^+^ debris. MNPs from both wild type and mutant animals showed engulfment of eGFP^+^ particles at 3 dpi, however fewer particles were internalized in *il34^-/-^* larvae compared to *il34^+/+^* siblings. Specifically, ∼57% (n=413) of the eGFP^+^ particles analyzed in *il34^+/+^* controls were associated with mCherry^+^ surfaces (≤10 nm depth), while ∼43% (n=307) had been internalized (≥20 nm depth). In *il34^-/-^* larvae, by comparison, 69% (n=602) of analyzed eGFP^+^ particles associated with mCherry^+^ surfaces, while only ∼31% (n=268) were internalized. The overall depth of particle internalization was also significantly decreased in *il34^-/-^* larvae compared to *il34^+/+^*siblings (Figure S5A; means=-0.08µm and-0.127µm, respectively). In addition to leukocyte mobilization, IL-34–CSF1R signaling also promotes MNP polarization to an anti-inflammatory (M2) phenotype^32,36^. Anti-inflammatory MNPs are highly phagocytic and representative of regeneration permissive tissue microenvironments. These data indicate that IL-34 signaling may drive MNP phagocytic function either directly, signaling through CSF1R, or indirectly due to the resultant delay in recruitment to the site of RPE damage.

**Figure 2.**
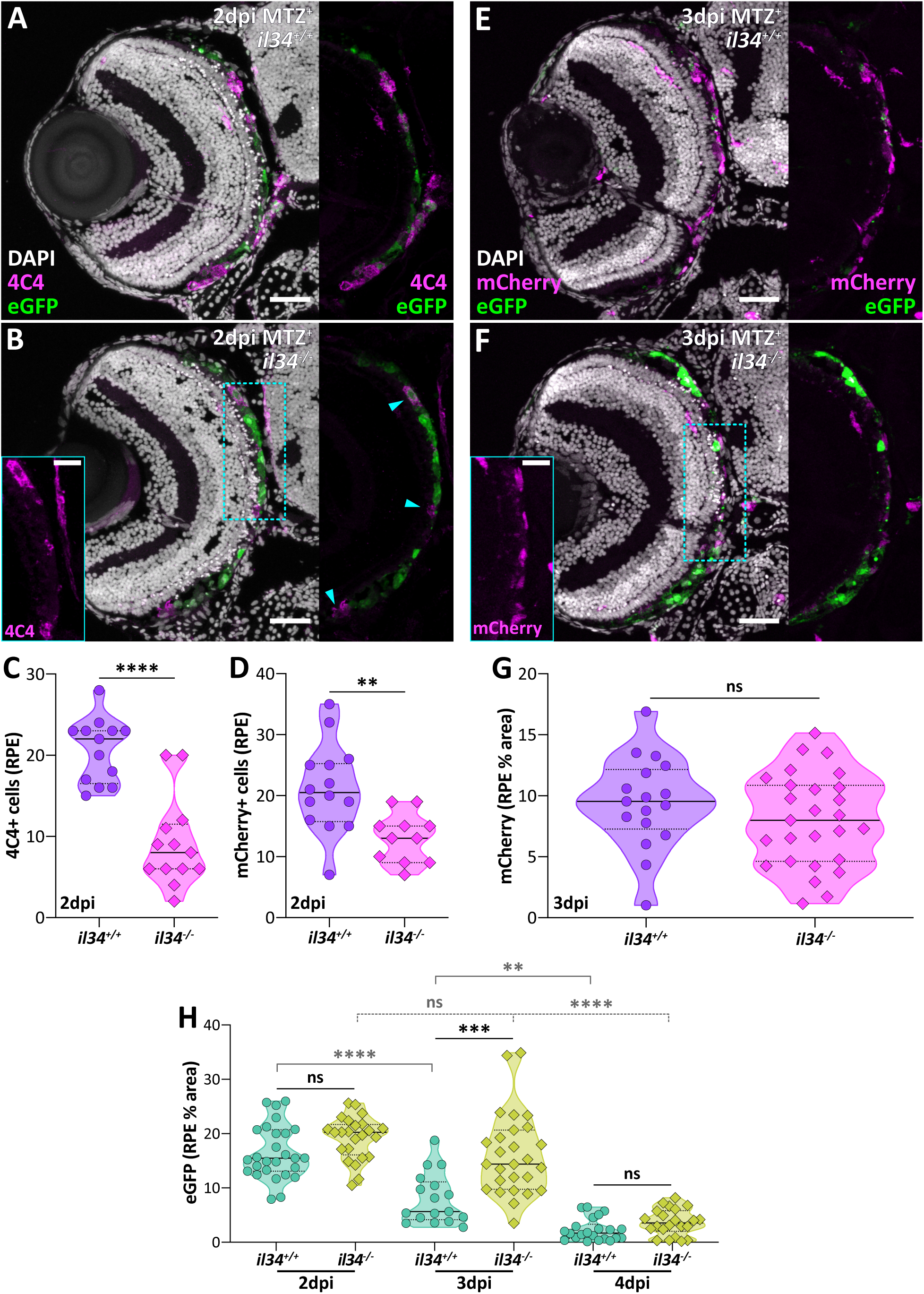
RPE-specific MNP recruitment is delayed in *il34* mutants. (A,B) Transverse confocal micrographs showing 4C4 *(magenta)* labeling in 2 dpi ablated (MTZ^+^) **(A)** *il34^+/+^* and **(B)** *il34^-/-^* larvae. **(B)** Digital zoom inset *(cyan outlines)* and *cyan arrowheads* highlight 4C4^+^ cells in the *il34^-/-^* animal. **(C,D)** Violin plots showing a significant decrease in **(C)** 4C4^+^ cells and **(D)** mCherry^+^ cells in the RPE of ablated (MTZ^+^) *il34^-/-^*larvae at 2 dpi. **(E,F)** Transverse confocal micrographs showing mCherry *(magenta)* labeling in 3 dpi ablated (MTZ^+^) **(E)** *il34^+/+^*and **(F)** *il34^-/-^* larvae. Digital zoom inset *(cyan outlines)* highlights mCherry^+^ cells in the *il34^-/-^*animal. **(G)** Violin plots showing no significant difference in RPE-localized mCherry expression (% area) between ablated (MTZ^+^) *il34^+/+^*and *il34^-/-^* larvae at 3 dpi. **(H)** Violin plots showing eGFP retention (% area) in ablated (MTZ^+^) *il34^+/+^* and *il34^-/-^* RPE at 2 dpi, 3 dpi, and 4 dpi. DAPI = nuclei *(white)*; *rpe65a:nfsB-eGFP* = RPE *(green)*; scale bars = 40µm; digital zoom scale bars = 20µm; **, p≤0.01; ***, p≤0.001; ****, p≤0.0001. Experimental replicates and statistical information can be found in Table S1.

In addition to eGFP^+^ debris, significant retention of pyknotic nuclei, a signature of apoptotic and necrotic cells, was also evident in *il34^-/-^* injury sites (Figure 2B,F). As RPE ablation also results in apoptosis in the photoreceptors that comprise the outer nuclear layer (ONL)^4^, we quantified cell death signatures in this collective functional unit. We detected significant increases in pyknotic nuclei at 2 dpi (Figure 3A–C) and TUNEL^+^ puncta at 3 dpi (Figure 3D–F) in *il34^-/-^* larvae, compared to wild type siblings. By 4 dpi, however, pyknotic nuclei counts were similar between *il34^+/+^*and *il34^-/-^* siblings (Figure 3G–I). These data align nicely with our eGFP results (Figure 2H) and support a model where the lag in MNP recruitment shifts the debris clearance window from 2 – 3 dpi to 3 – 4 dpi in *il34* mutants. As we did not detect significantly different eGFP or pyknosis levels in *il34^-/-^* animals at 4 dpi (Figures 2H and 3G–I, respectively), the residual debris present at this timepoint is a likely combination of pigment-containing phagocytic MNPs and immature but regenerating peripheral RPE tissue undergoing differentiation and melanogenesis (Figure 1D). Importantly, there was no difference in cell death indicators between unablated *il34* wild type or mutant groups at 6 dpf (Figure S3C), 8 dpf (Figure 3F), or 9 dpf (Figure 3I). Baseline numbers of pyknotic nuclei were slightly elevated in unablated *il34^-/-^* retinae at 7 dpf (Figure 3C), but this may represent a transient peak seen in the ONL during normal retinal neurogenesis^37^. IL-34 signaling has been shown to drive proliferation, however characterization of *il34* mutants by Kuil *et al.*^23^ showed that loss of *il34* does not appear to influence the proliferative capacity of MNPs. Accordingly, we did not detect any differences in proliferation dynamics between *il34^+/+^* and *il34^-/-^* RPE injury sites (Figure S6), which include recruited MNPs (Figure 2)^8^. We previously characterized a ∼48-hour window during which MNPs infiltrate the RPE injury site (2 dpi), perform pro-regenerative actions, and then leave (4 dpi)^8^. Fitting within this timeline, these data show that IL-34 signaling mediates precisely timed MNP recruitment at 2 dpi, which initiates subsequent cellular debris removal and progression of RPE regeneration. As *il34* is expressed by damaged RPE, these findings strongly support a model in which IL-34 signaling originates in the RPE to activate MNP-driven tissue repair mechanisms.

**Figure 3.**
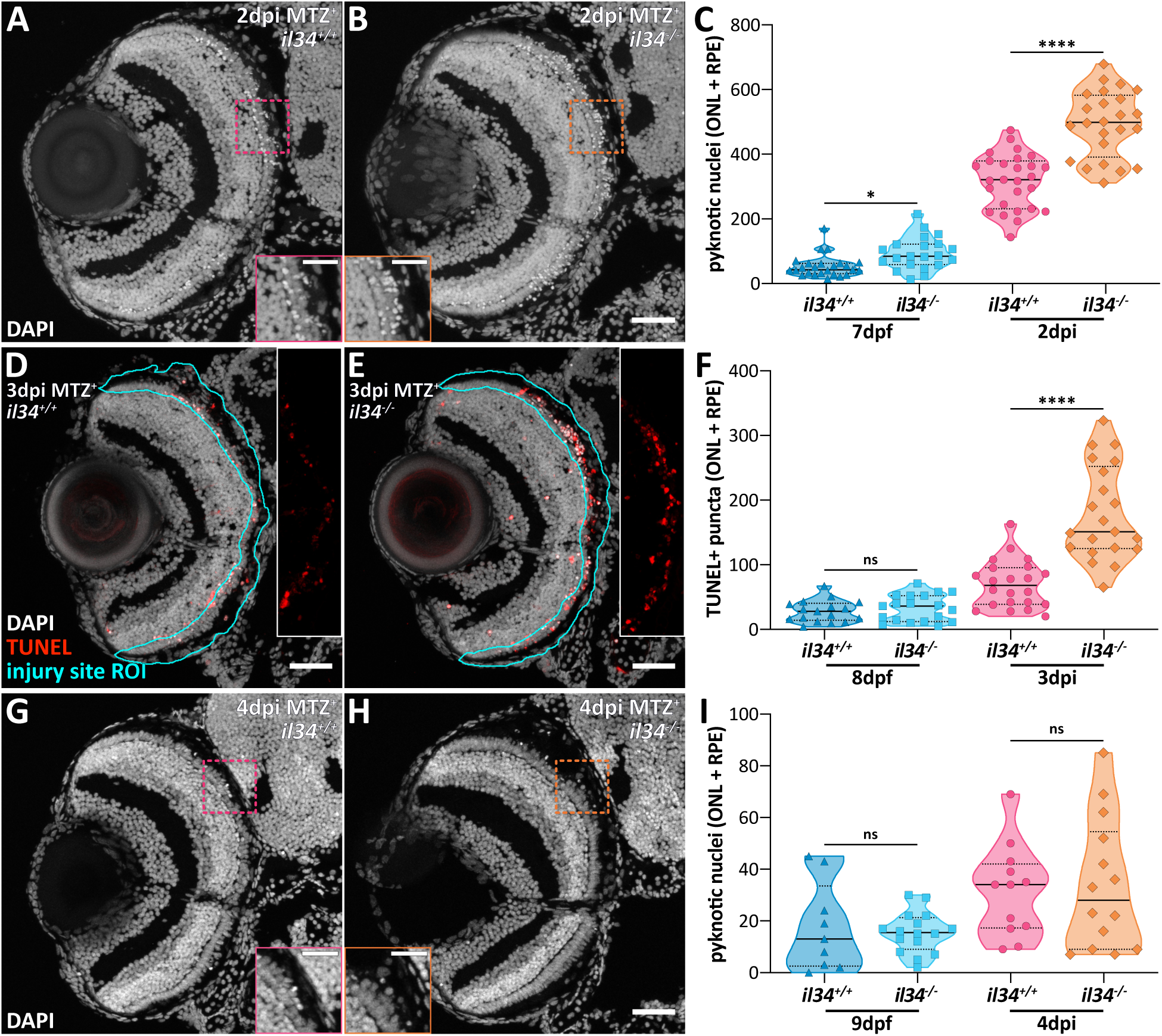
*il34* mutant RPE retain apoptotic/necrotic cells at early timepoints post-injury. (A,B) Transverse confocal micrographs showing DAPI counterstaining *(white)* in 2 dpi ablated (MTZ^+^) **(A)** *il34^+/+^*and **(B)** *il34^-/-^* larvae. Digital zoom insets *(orange and pink outlines)* emphasize increased numbers of pyknotic nuclei in the *il34^-/-^* animal. **(C)** Violin plots showing pyknotic nuclei counts in 7 dpf unablated (MTZ^-^) and 2 dpi ablated (MTZ^+^) larvae. **(D,E)** Transverse confocal micrographs showing TUNEL labeling *(red)* in 3 dpi ablated (MTZ^+^) **(D)** *il34^+/+^* and **(E)** *il34^-/-^* larvae. *Cyan outlines* designate regions of interest (ROI) encompassing the photoreceptors (ONL) and RPE. Single-channel *insets* (no zoom) highlight increased numbers of TUNEL^+^ puncta in the *il34^-/-^* animal. **(F)** Violin plots showing quantification of TUNEL in 8 dpf unablated (MTZ^-^) and 3 dpi ablated (MTZ^+^) larvae. **(G,H)** Transverse confocal micrographs showing DAPI counterstaining *(white)* in 4 dpi ablated (MTZ^+^) **(G)** *il34^+/+^*and **(H)** *il34^-/-^* larvae. Digital zoom insets *(orange and pink outlines)* emphasize minimal pyknotic nuclei in both animals. **(I)** Violin plots showing pyknotic nuclei counts in 9 dpf unablated (MTZ^-^) and 4 dpi ablated (MTZ^+^) larvae. Scale bars represent **(B,D,E,H)** 40µm and (**A,B,G,H** digital zooms) 20µm; *, p≤0.05; ****, p≤0.0001. Experimental replicates and statistical information can be found in Table S1.

### The RPE regenerates normally in late-stage *il34* mutant larvae despite early MNP recruitment delays

Our prior characterization showed that RPE and underlying photoreceptor regeneration is largely complete by 14 dpi^4^. At this timepoint, late-stage larvae featured mature markers across the entire RPE layer, including expression of *rpe65a*-driven eGFP, restoration of epithelial characteristics (e.g., apical-basal polarity), and mature melanosomes^4^. Perhaps owing to the highly specialized function and comparatively late-phase neurogenesis of cells in the ONL^37^, photoreceptors showed some residual central disorganization at 14 dpi that was remedied at later ages^4^. Despite delays in MNP recruitment and debris clearance, *il34^-/-^* RPE regenerated by 14 dpi (Figure 4). Consistent with larval results, unablated 19 dpf age-matched *il34^+/+^*and *il34^-/-^* siblings showed typical central retinal morphologies, including elongated photoreceptor outer segments (Figure 4B; *dotted yellow lines*) and polarized RPE with apical microvilli (Figure 4B; *cyan arrowheads*). At 14 dpi, a third of *il34^+/+^* (2/6) and 5 out of 6 *il34^-/-^*retinae showed central RPE and ONL morphologies similar to wild type and mutant unablated controls (Figure 4C). The remaining animals (4/6 *il34^+/+^* and 1/6 *il34^-/-^*) had morphologically normal RPE, although some central outer segments remained disorganized and/or appeared truncated (Figure 4D; *arrowheads*), as expected^4^. Nevertheless, these animals displayed prominent interdigitation of outer segments with RPE apical microvilli (Figure 4E), demonstrating successful reestablishment of the photoreceptor–RPE functional unit. These findings indicate that there are likely multiple pro-regenerative signals converging to activate MNPs; indeed, pathway synergy has been reported in other complex tissue regeneration models, like the zebrafish retina^38–42^.

**Figure 4.**
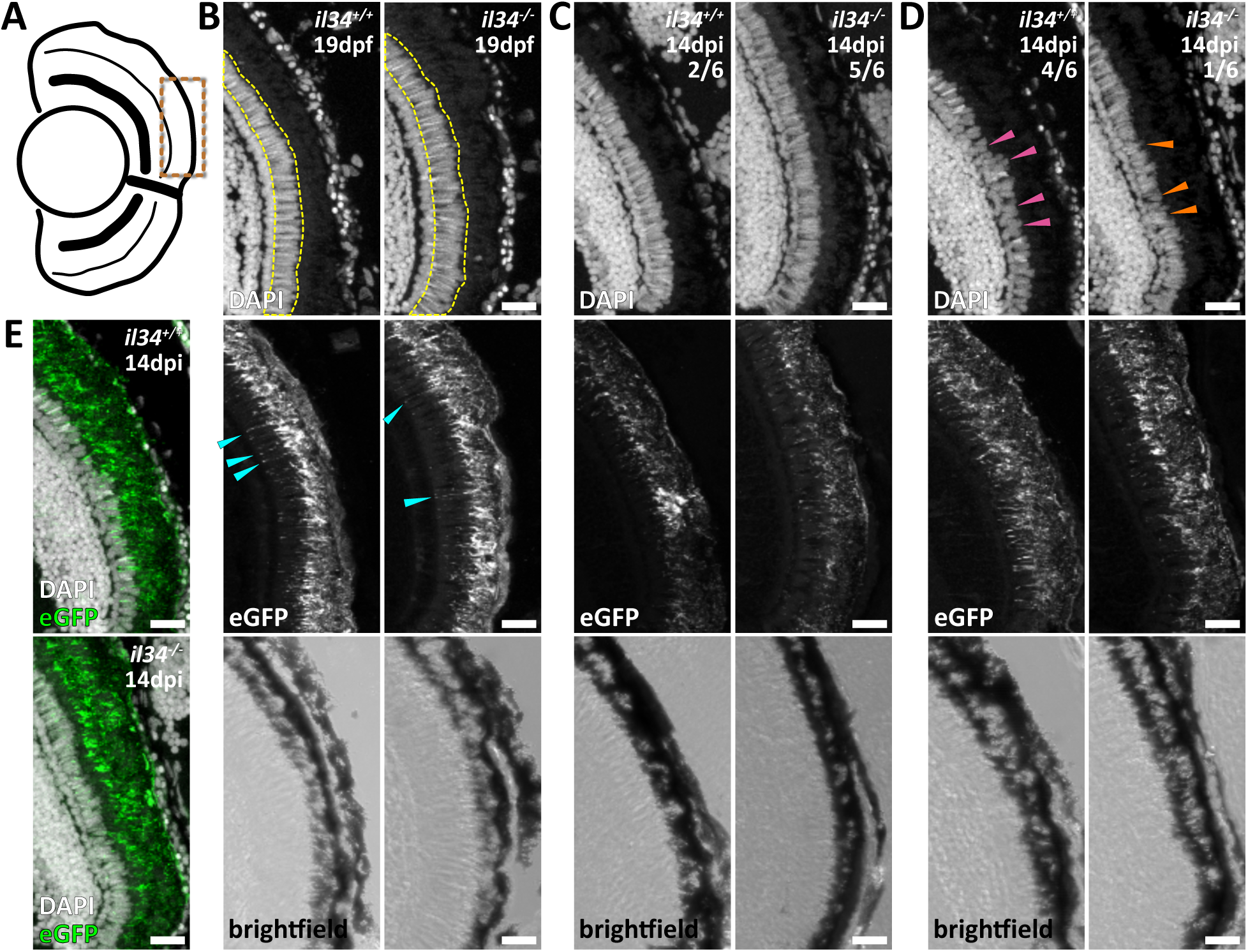
*il34* mutant RPE regenerate normally by 14 dpi despite early timepoint deficiencies. **(A)** Schematic of a late-stage larval eye depicting the central-dorsal region zoomed in on in **B–E** *(bronze dotted line)*. **(B,C)** Transverse confocal micrograph digital zooms showing **(B)** 19 dpf unablated (MTZ^-^) *il34^+/+^* and *il34^-/-^* larvae and **(C)** 14 dpi ablated (MTZ^+^) *il34^+/+^* and *il34^-/-^* animals with no discernible phenotype. Examples of uniformly normal photoreceptors (elongated outer segments) and RPE (apical microvilli) morphologies are highlighted with *yellow dotted lines* and *cyan arrowheads*, respectively. **(D)** Digital zooms showing 14 dpi ablated (MTZ^+^) *il34^+/+^* and *il34^-/-^* larvae with normal RPE but truncated or disorganized outer segments *(arrowheads)*. **(E)** DAPI and eGFP overlays showing the same retinae as in **(D)**. All scale bars = 20µm; DAPI = nuclei *(white)*; *rpe65a:nfsB-eGFP* = RPE *(white and green)*. Experimental replicate information can be found in Table S1.

### *csf1ra/b* double mutants show severe tissue degeneration phenotypes at peak regenerative timepoints

One of the IL-34 receptors, CSF1R, is required for the function and embryonic tissue dispersion of MNPs in vertebrates^24,31,43–47^. Zebrafish harboring loss-of-function mutations in both *csf1ra* and *csf1rb* (herein, *csf1r^DM^*), which, together share functional homology with mammalian CSF1R^24^, lack microglia almost completely and show significant systemic deficiencies in some MNP populations, but likely retain circulating monocytes^24,43,48–50^. Given this is a microglia deficient model for which we utilized only lgals3bpb/4C4 staining, we will reference microglia specifically throughout the discussion of CSF1R-related results. Leveraging this unique model as a tool to eliminate not only the downstream effects of IL-34–CSF1R signaling, but all the reparative roles microglia play during RPE regeneration, we crossed our *Tg(rpe65a:nfsB-eGFP)* line to *csf1ra/b* mutants. Consistently, we observed almost no 4C4^+^ cells in unablated *csf1r^DM^* eyes at 8 dpf, when compared to *csf1r^+/+^* siblings (Figure S4G–I). This was similar after ablation as well, and we counted zero or very few 4C4^+^ cells in *csf1r^DM^* RPE at 3 dpi, whereas infiltration was robust in *csf1r^+/+^* siblings (Figure 5A–C), as expected. As with the *il34* mutants, close examination of ablated *csf1r^DM^* eyes revealed retention of eGFP^+^ (Figure 5b’’) and pyknotic (Figure 5b’’’) debris, which were significantly elevated compared to sibling controls (Figure 5D,E). Pearson correlation analyses showed a strong positive relationship between eGFP and pyknotic nuclei (p<0.0001, r=0.78), and a very strong negative relationship between the number of 4C4^+^ cells and both eGFP (p<0.0001, r=-0.80) and pyknotic nuclei (p<0.0001, r=-0.83) (Figure S7), further supporting a central role for microglia in injury site debris clearance. Ablated *csf1r^DM^* retinae showed additional central tissue regeneration deficiencies that persisted at peak regeneration (4 dpi), including severe zpr-1^+^ cone photoreceptor degeneration phenotypes, compared to *csf1r^+/+^* sibling controls (Figure 5F,G). We previously showed that degeneration of the underlying photoreceptors is a secondary consequence of RPE ablation in our model, but that photoreceptors were visibly recovering by 4 dpi in a wild type genetic background (e.g., Figure 5F)^4^. *csf1r^DM^* larvae, however, showed severely truncated or missing central photoreceptor tissue in addition to continued retention of eGFP^+^ debris at 4 dpi (Figure 5G). RPE and photoreceptor phenotypes were collectively used to determine the central-most limits of regenerating tissue, and dorsal and ventral angle measurements made based on these edges (e.g., Figure 5F,G; *arrowheads*) showed a significant decrease in tissue recovery in the 4 dpi *csf1r^DM^* cohort (Figure 5H). Like *il34* mutants, loss of *csf1ra/b* did not yield any detectable ocular developmental or maintenance phenotypes, as unablated RPE morphologies (Figure S2E–H) and initial ablation severities (Figure S3D–F) did not differ between *csf1r^+/+^* and *csf1r^DM^* larvae. Together, these findings show that microglia are dispensable for normal RPE development but perform an essential debris clearing function that is required to advance tissue regeneration post-RPE injury.

**Figure 5.**
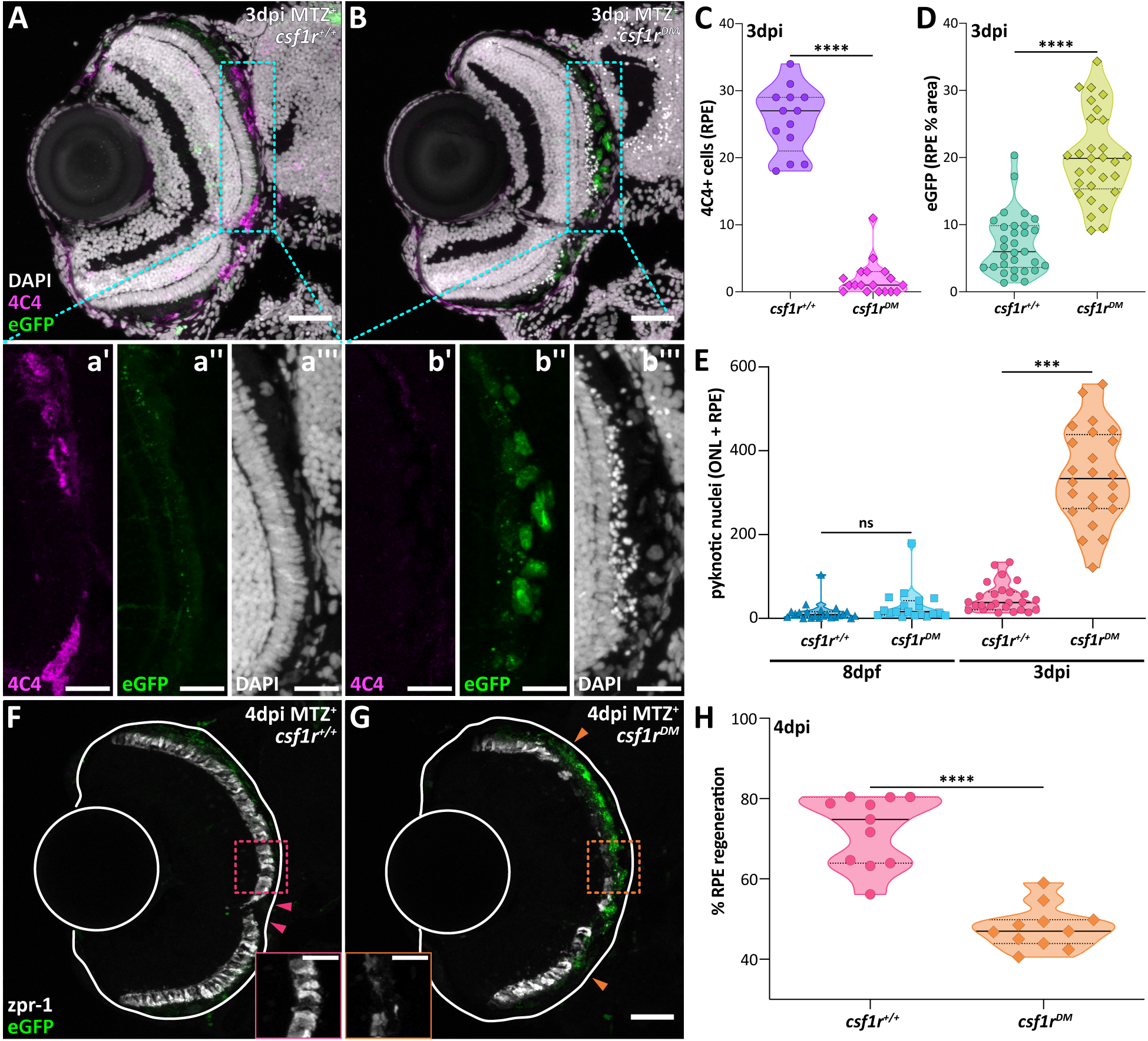
Tissue regeneration is severely impaired in *csf1ra/b* double mutants, which lack microglia. (A,B) Transverse confocal micrographs showing 4C4 *(magenta)* labeling in 3 dpi ablated (MTZ^+^) **(A)** *csf1r^+/+^* and **(B)** *csf1r^DM^* larvae. **(a’–b”’)** Single-channel digital zooms *(cyan outlines)* highlight the **(b’)** lack of 4C4^+^ cells *(magenta)*, and accumulation of **(b”)** ablated eGFP^+^ *(green)* and **(b”’)** pyknotic *(white)* debris in the *csf1r^DM^* animal. **(C)** Violin plots showing 4C4^+^ cell counts in ablated (MTZ^+^) *csf1r^+/+^* and *csf1r^DM^* larvae at 3 dpi. **(D,E)** Violin plots showing **(D)** eGFP^+^ signal (% area) and **(E)** pyknotic nuclei counts in ablated (MTZ^+^) *csf1r^+/+^* and *csf1r^DM^* larvae at 3 dpi. **(F,G)** Transverse confocal micrographs showing expression of the cone photoreceptor marker, zpr-1 *(white)*, and the RPE *(green)* in 4 dpi ablated (MTZ^+^) **(F)** *csf1r^+/+^* and **(G)** *csf1r^DM^* larvae. *Solid white lines* outline the lens and outer limits (sclera) of the eye. *Arrowheads* designate the central-most limits of regenerated tissue. **(H)** Violin plots showing percent RPE regeneration in ablated (MTZ^+^) *csf1r^+/+^*and *csf1r^DM^* larvae at 4 dpi. Scale bars = 40µm; digital zoom scale bars = 20µm; ***, p≤0.001; ****, p≤0.0001. Experimental replicates and statistical information can be found in Table S1.

### The RPE fails to regenerate in late-stage *csf1ra/b* double mutant larvae

Given the severity of lingering tissue degeneration in 4 dpi *csf1r^DM^* larvae (Figure 5G), we aged *csf1ra/b* animals to 14 dpi to determine long-term RPE regeneration success. As retinal neurons and RPE cells are continuously born from the peripheral eye in teleosts^51–53^, we focused analyses on the central-most tissues of the ONL and RPE, adjacent to the optic nerve (e.g., Figure 6B), which was part of the original ablation site induced at 5 – 6 dpf. We implemented a scoring system to assess phenotypes across 14 dpi *csf1r^+/+^* and *csf1r^DM^* late-stage larvae (Figure 6A). Ablated animals with no phenotype (“none”) were indistinguishable from 19 dpf unablated controls, which showed elongated photoreceptor outer segments that aligned parallel to and interdigitated with the RPE apical microvilli regardless of genotype (Figure 6C). Only *csf1r^+/+^*retinae (4/12) displayed phenotypes analogous to unablated controls (Figure 6A,D). Mild phenotypes were assigned to retinae displaying morphologically normal RPE but minimal outer segment disorganization or truncation, much like the *il34* cohorts (Figure 4D) and previously characterized 14 dpi larvae^4^. Again, only *csf1r^+/+^* retinae (4/12) displayed mild phenotypes (Figure 6A,E). Both *csf1r^+/+^* and *csf1r^DM^* retinae exhibited moderate central phenotypes (Figure 6A,F; 4/12 and 2/6, respectively) characterized by areas of lost or severely truncated outer segments (e.g., Figure 6F; *magenta arrowheads*), loss of RPE apical microvilli/polarity (e.g., Figure 6F; *green arrowhead*), and/or retention of pyknotic nuclei (e.g., Figure 6F; *white arrows*). Lastly, the majority of *csf1r^DM^* larvae (4/6) exclusively presented with severe phenotypes (Figure 6A,G). Hallmarks of these retinae were heavily pigmented but disorganized central tissue clumps that lacked eGFP expression (e.g., Figure 6G; *asterisks*), some of which obstructed the RPE monolayer, preventing polarization of under-or overlying eGFP^+^ RPE tissue (e.g., Figure 6G; *green arrowheads*). Beyond the absence of eGFP expression, it is unlikely that these pigmented tissue occlusions represent mature and/or functional RPE as the immediately underlying photoreceptor layer appeared hypotrophic and lacked outer segments in some areas (e.g., Figure 6G; *magenta arrowheads*). Figure 6A summarizes all scored 14 dpi larvae and emphasizes the predominance of normal or mild phenotypes in *csf1r^+/+^* animals, while the majority of the *csf1r^DM^* cohort phenotypes were severely impaired. The centrality and prominent pigmentation of residual occlusions in severe 14 dpi *csf1r^DM^* retinae coupled with the degeneration of the underlying ONL strongly suggest an ablated RPE origin that is, based on poor tissue health at 3 – 4 dpi (Figure 5), a result of remnant, uncleared debris in the RPE injury site. The ultimate failure of *csf1r^DM^* animals to regenerate in the long term is striking given the robust reparative capacity of the zebrafish model system. Collectively, these data show that the active role of microglia as phagocytes is indispensable for RPE and photoreceptor regeneration.

**Figure 6.**
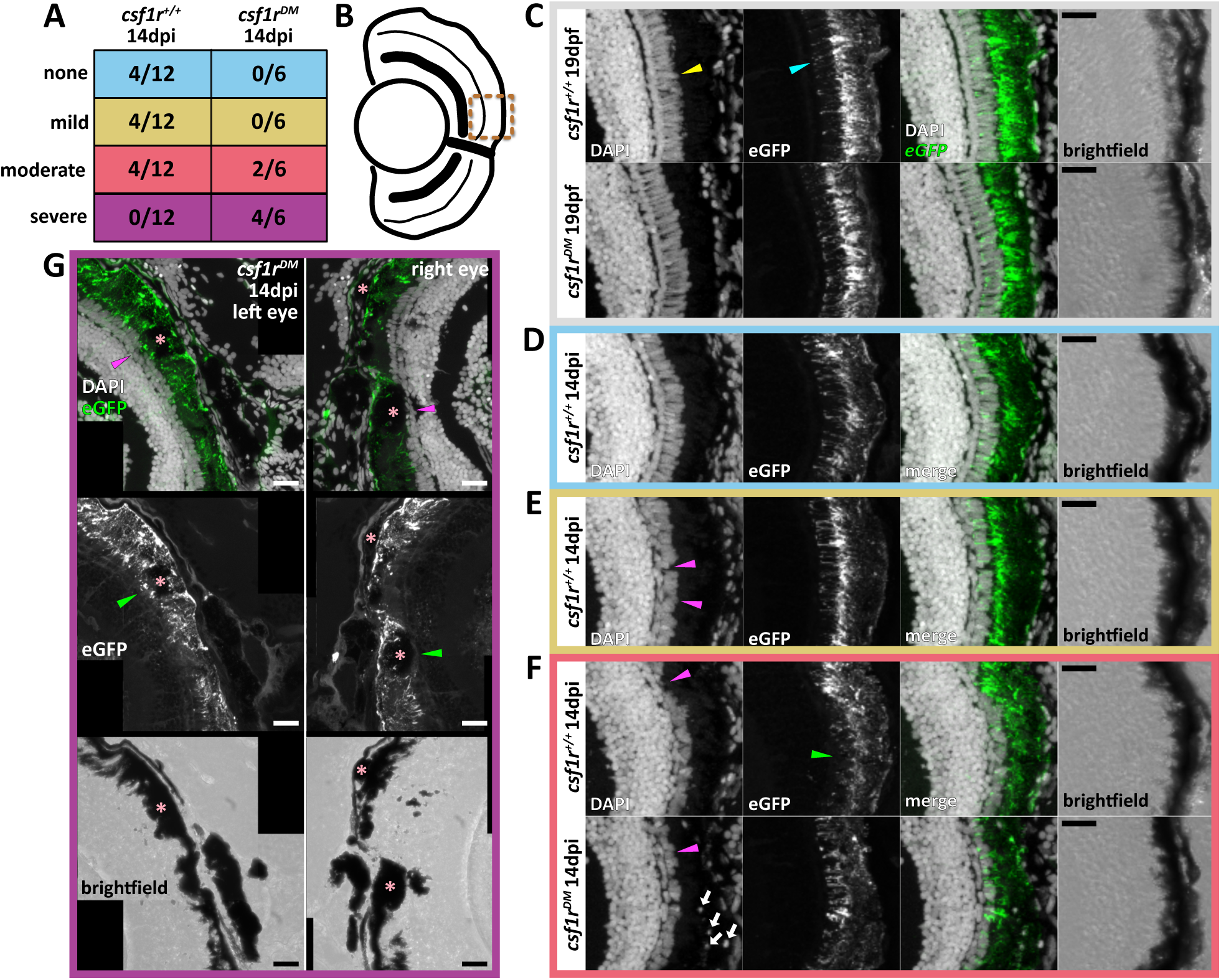
*csf1ra/b* double mutant zebrafish show impaired RPE regeneration phenotypes at 14 dpi. **(A)** Table showing the degree of phenotype severity in 14 dpi larvae *(n=12 total, csf1r^+/+^; n=6 total, csf1r^DM^)*. **(B)** Schematic of a late-stage larval eye depicting the central-dorsal region zoomed in on in **C–F** *(bronze dotted line)*. **(C)** Transverse confocal micrograph digital zooms showing 19 dpf unablated (MTZ^-^) *csf1r^+/+^*and *csf1r^DM^* larvae. Examples of uniformly normal photoreceptor (elongated outer segments) and RPE (apical microvilli) morphologies are highlighted with *yellow* and *cyan arrowheads*, respectively. **(D–F)** Transverse confocal micrograph digital zooms showing 14 dpi ablated (MTZ^+^) *csf1r^+/+^* or *csf1r^DM^*larvae with **(D)** no discernible phenotype, **(E)** mild PR and RPE phenotypes, and **(F)** moderate PR and RPE phenotypes. *Magenta arrowheads* point out areas of sparse, absent, or hypertrophic outer segments. *Green arrowhead* indicates an area where apical microvilli are missing or truncated. *White arrows* indicate cellular debris retention. **(G)** Stitched transverse confocal micrographs showing the right and left eyes of a 14 dpi ablated (MTZ^+^) *csf1r^DM^* larva with severe photoreceptor and RPE phenotypes. *Green* and *magenta arrowheads* point to areas of abnormal RPE and photoreceptor outer segments, respectively, that localize with large, pigmented debris deposits *(asterisks)*. All scale bars = 20µm; DAPI = nuclei *(white)*; *rpe65a:nfsB-eGFP* = RPE *(white and green)*. Experimental replicate information can be found in Table S1.

## Discussion

In an effort to dampen the global burden that RPE degenerative diseases, like AMD, impose on the healthcare industry and patient quality of life, the last ∼15 years has seen a remarkable push to develop stem cell-based RPE replacement therapies for clinical application^54^. However, due to complex etiologies that encompass genetic and/or environmental factors, coupled with the limited regenerative capacity of mammalian tissue, there are still no cures for RPE degenerative diseases. Further, despite a massive effort in the clinical realm, we know little about the signals driving intrinsic RPE regeneration. Our findings here expand the field by validating novel regenerative mechanisms – IL-34 signaling and MNP-mediated debris clearance – that regulate intrinsic RPE repair. Building upon previous studies^18,30^, we show that *il34* is required for precise temporal recruitment of MNPs (Figure 7B) and that MNP injury site localization is required for debris clearance (Figure 7B,C). Injury site debris clearance is a critical step that drives resolution of inflammation and is essential for tissue repair in both regeneration-competent^55–57^ and-incompetent (mammalian)^18,58,59^ species. The zebrafish lines used here encompass impairment severities ranging from (at most) a 24-hour MNP recruitment delay (*il34* mutants) to a near total loss of microglia (*csf1r^DM^*); and our results illuminate respective parallels in tissue regeneration severity, from a short-term lapse in debris clearance to a long-term failure to regenerate RPE and ONL tissues (Figure 7B,C). Together, our findings underscore the critical requirement for debris removal by MNPs after RPE damage for regeneration to occur.

**Figure 7.**
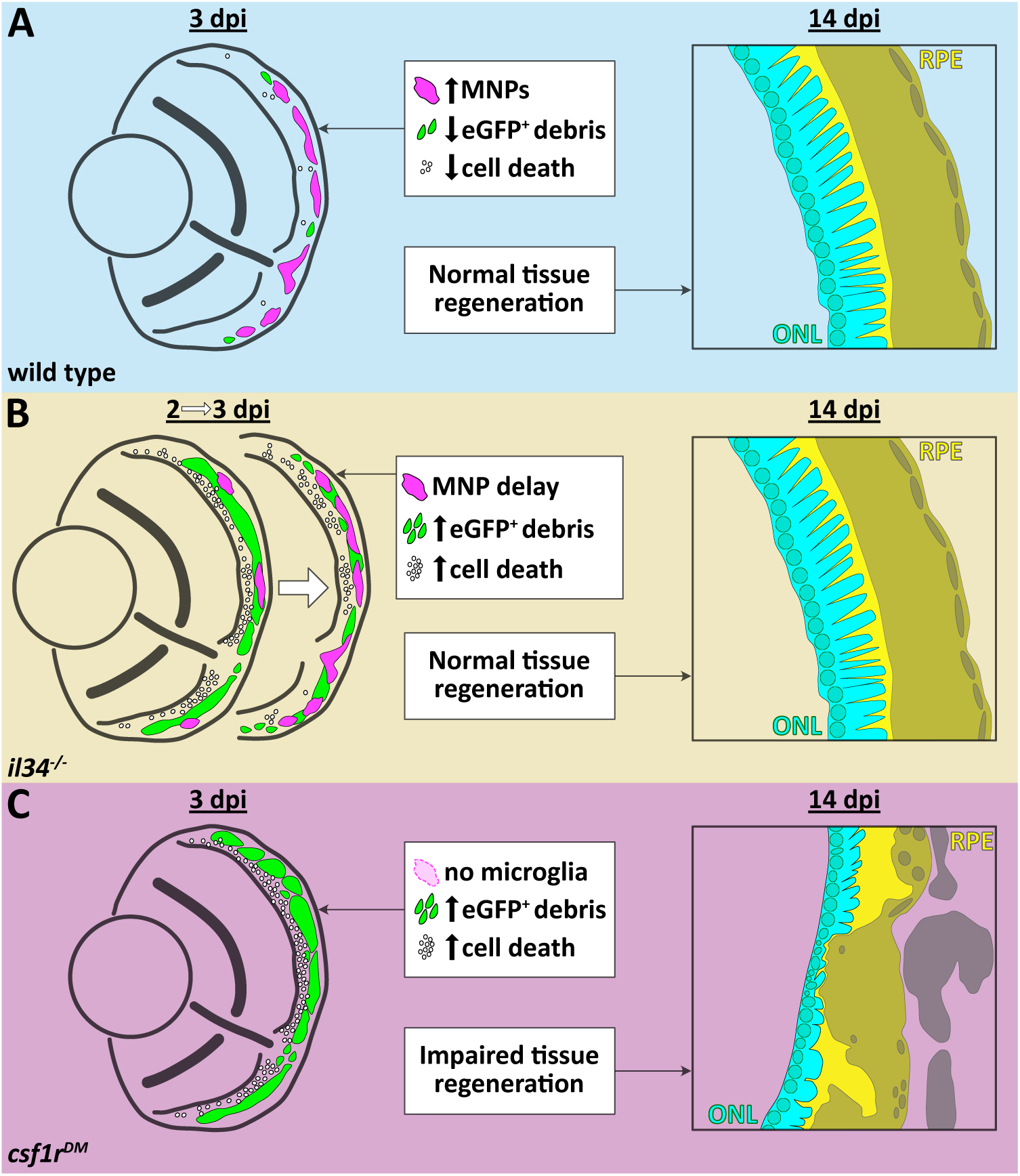
The consequences of *il34* or *csf1ra/b* loss on RPE regeneration. **(A)** Schematic representation of wild type larval eyes at 3 dpi and 14 dpi. MNPs *(magenta)* are recruited to the RPE layer, eGFP^+^ debris *(green)* and cell death markers *(white)* are low, and ONL and RPE tissues regenerate normally. **(B)** Schematic representation of *il34^-/-^*larval eyes at 2 – 3 dpi and 14 dpi. MNP *(magenta)* recruitment is delayed at 2 dpi but recovers by 3 dpi; eGFP^+^ debris *(green)* and cell death markers *(white)* accumulate; however, ONL and RPE tissues regenerate normally. **(C)** Schematic representation of *csf1r^DM^* larval eyes at 3 dpi and 14 dpi. Microglia *(magenta)* are absent, there is heavy accumulation of eGFP^+^ debris *(green)* and cell death markers *(white)*, and ONL and RPE tissues fail to regenerate.

While MNPs can play a reparative role, prolonged accumulation and immune activation can lead to collateral tissue death and degenerative disease progression if unresolved, as is the case in AMD^60^. There is also mounting evidence that distinct MNP populations differentially contribute to ocular pathology^61^. Specifically, monocyte-derived macrophages have been linked to AMD pathogenesis^62^, whereas microglia have been associated with protection against retinal degeneration^18,62–64^. Our study provides insight into this paradox. As established by others, and corroborated here, *csf1r^DM^* zebrafish lack nearly all microglia in the retina and brain^24,43,48,49^; however, findings from optic nerve^49^ and spinal cord^50^ regeneration studies indicate that *csf1r^DM^* animals retain circulating monocyte populations, which are capable of differentiating into macrophages in response to tissue injury. For example, Tsarouchas *et al.*^50^ showed that microglia-deficient *csf1r^DM^* zebrafish successfully repaired spinal cord transection lesions after robust infiltration of mfap4-labeled cells (e.g., monocytes/macrophages^65^), indicating that recruitment of peripheral macrophages, not microglia, is required for regeneration in this model. Conversely, we found that *csf1r^DM^*animals showed a complete failure to regenerate RPE and photoreceptor cells by 14 dpi, which is noteworthy given the robust reparative potential of the zebrafish and the likelihood that these animals harbor injury-responsive monocyte-derived macrophage populations. Although we did not examine mfap4 expression, our results make a strong case for microglia as the predominant leukocyte required for successful RPE regeneration and indicate that different MNP populations may also play functionally distinct roles during regenerative responses. Fitting a perspective that has been proposed within the context of retinal degeneration and AMD^61^, our findings are significant in suggesting that differential modulation of microglia and/or monocyte-derived macrophages may be needed to achieve pro-regenerative outcomes during RPE degenerative disease intervention. Similar to zebrafish, microglia-deficient *Csf1r^-/-^* mice also retain normal levels of blood-derived monocytes^46^ and future comparative studies using these vertebrate models may be useful to parse apart the distinct roles of different MNP populations during RPE damage and repair responses.

Similarly, while we will not make the argument that either RPE-ablated mutant line used herein is an AMD model, *csf1r^DM^* late-stage larvae display phenotypic similarities that hint at underlying AMD-relevant biological processes that would be interesting to study in isolation from protective microglia responses. For example, the debris existing between the RPE and photoreceptors and basal to the RPE in *csf1r^DM^* retinae (e.g., Figure 6G; *asterisks*) is spatially and morphologically similar to drusenoid deposits in late-stage atrophic AMD where, like in our model, RPE and photoreceptor tissue is lost and/or highly disorganized^66^. Thus, RPE-ablated *csf1r^DM^* zebrafish may be useful for studying the consequences of debris size and localization without continued clearance. As we assessed only lgals3bpb/4C4 localization, it remains to be determined if prolonged inflammatory input from monocyte-derived macrophages and/or residual damaged tissue also contributes to sustained RPE and ONL degeneration in *csf1r^DM^* retinae. Additionally, aberrant RPE and photoreceptor cell death is a prominent early phenotype in both *il34* and *csf1ra/b* mutants and is also a hallmark of late-stage atrophic AMD. Interestingly, in *csf1r^DM^*animals, disorganized tissue footprints remain regionally contained to the central RPE/retina, at least at 14 dpi, as there is clear delimitation between healthy and unregenerated tissues. In atrophic AMD, lesions expand with disease progression, and part of this spread is likely due to the pathogenic role of MNPs^60^. Thus, RPE-ablated *csf1r^DM^* zebrafish may also be a useful model to identify cell–cell signaling mechanisms at the interface of healthy and damaged tissues that contribute to the restriction or spread of tissue degeneration independent of protective immune responses.

While this study utilizes a systemic *il34* mutant zebrafish line, our results support a model where IL-34 derived from damaged RPE signals to MNPs to initiate pro-regenerative functions. Like many cytokine–receptor pathways, IL-34 signaling is multifaceted and complex. For example, autocrine/paracrine IL-34–CSF1R signaling functions upstream of critical MNP effector activities^15–17^, as mentioned, and in leukocyte-independent contexts as well^20^. IL-34 is also a putative ligand for several other receptors, including syndecan-1 (CD138)^13^, receptor-type protein-tyrosine phosphatase zeta (PTP-ζ)^12^, and trigger receptor expressed myeloid 2 (TREM2)^14^. Syndecan-1 and TREM2, like CSF1R, are predominately linked to MNP functions, whereas PTP-ζ is expressed by mature cells and progenitors of the CNS^67^. We did not previously detect upregulation of *il34* in mCherry^+^ MNPs responding to RPE ablation, or appreciable levels of *csf1ra/b* or *ptprz1a/b* receptor genes in RPE at early (2 dpi) or peak (4 dpi) regenerative timepoints by RNA-seq^8^; thus, it is unlikely that autocrine IL-34 signaling mechanisms are at play in our model. Given the neuroprotective role of IL-34 signaling^18–20^, and the prominent cell death phenotypes detected in the ONL, we cannot rule out the possibility that alternate paracrine IL-34 signaling mechanisms (e.g., RPE–ONL) also impact cell survival in synergy with MNP recruitment and debris clearance. Nevertheless, our findings support a pro-regenerative paracrine RPE–MNP IL-34 signaling mechanism, which is exemplified by the RPE-specific expression of *il34*, the MNP recruitment delay from 2 to 3 dpi and the concomitant lapse in appreciable debris clearance from 3 to 4 dpi in *il34* mutants. These results have potential translational relevance in the context of RPE degenerative disease intervention as IL-34-dependent microglia have been shown to be RPE-protective in a mouse model of retinal degeneration^18^, pointing to mechanistic conservation between teleosts and mammals.

Ultimately, despite aberrant cell death and debris clearance phenotypes, MNP influx normalized by 3 dpi and RPE regeneration was successful in *il34* mutants in the long term (Figure 7B). This indicates that other signals, likely other cytokines/chemokines, function to overcome the loss of *il34*, which is highly plausible as acute injuries evoke strong and complex inflammatory responses in zebrafish^5^. For example, the chemokine, CXCL8, and matrix metalloproteases, MMP9 and MMP13, have been implicated in leukocyte recruitment during heart regeneration^68^. Indeed, we showed that damaged and/or regenerating RPE upregulate expression of *cxcl8a*, *mmp9,* and *mmp13b*^8^, suggesting this could be an IL-34-independent mechanism for MNP recruitment in our model as well. It should be noted, however, that the significant delay in MNP recruitment with just the loss of *il34* highlights the importance of this cytokine for RPE regeneration. Further, RPE debris removal in *il34* mutants at 14 dpi suggests other signals may also influence MNP phagocytic functions. Interleukin-11 (IL-11) is a potential candidate signal as *il11a* and *il11b* are upregulated in regenerating zebrafish RPE^8^ and IL-11 signaling modulates a multitude of leukocyte effector activities, including phagocytosis and other anti-inflammatory actions^69–71^. Thus, it is likely that multiple signals converge to enhance MNP recruitment and other pro-regenerative functions, and the intricacies of cytokine/chemokine pathway cooperation remain to be examined.

In summary, this study shows that IL-34 signaling and MNP-mediated injury site debris clearance mechanisms drive intrinsic RPE regeneration. Our conclusions expand knowledge in the field by pinpointing RPE-derived IL-34 as a critical signal for MNP recruitment and microglia as the predominant leukocytes promoting RPE repair. Further, we identify a model of complete RPE regeneration failure, which provides a unique, multipurpose *in vivo* platform that could be utilized to understand underlying degenerative disease-related RPE pathologies and evaluate the distinct roles of different MNP populations in RPE reparative responses.

## Materials and Methods

### Zebrafish lines, maintenance, and genotyping

Zebrafish (see Table S2 for line information) were handled according to The University of Texas at Austin Institutional Animal Care and Use Committee (IACUC) regulations. Adult zebrafish were housed in a recirculating facility (Tecniplast, Buguggiate VA, Italy) at 28.5 °C on a 14-/10-hour light/dark cycle. Adults were fed a standard diet of ZEBRAFEED (400 – 600 µm) by SPAROS (Tecniplast). Embryos and larvae were kept in a 28.5 °C incubator (no light/dark cycle) until euthanasia by tricaine (MS-222; Pentair Aquatic EcoSystems, Inc., London, United Kingdom) overdose. At the time of screening (5 dpf), eGFP^+^ siblings were randomly assigned to unablated or ablated experimental groups. Larvae euthanized ≤9 days post-fertilization (dpf) were unfed^72^. Larvae aged to 19 dpf were put into rotifer (S type; Reed Mariculture, Campbell, CA, USA) polyculture at 7 dpf for continuous feeding until euthanasia. Larval sex differences were not a factor in this study as zebrafish do not become sexually mature/dimorphic until 2.5 – 3 months of age. Genotyping was performed on any of the following 96-well thermal cyclers: T100, C1000 Touch, or PTC-100 (all Bio-Rad, Hercules, CA, USA). Zebrafish from *il34^re^*^03^ crosses were genotyped and PCR products were subsequently digested with HpyCH4IV (New England Biolabs, Ipswich, MA, USA) to distinguish wild type and mutant alleles. Genotyping PCR products from *csf1ra^j4e^*^1^ and *csf1rb^re^*^01^ animals were digested with SpeI or MspI (New England Biolabs), respectively. Uncut PCR products from all lines were periodically sequenced to confirm wild type and mutant genotyping methodologies. Genotyping primer sequences and additional information can be found in Table S2.

### RPE ablation using metronidazole (MTZ)

Ablation methodology was performed as described^30^. Briefly, embryos from *rpe65a:nfsB-eGFP* x AB outcrosses, *rpe65a:nfsB-eGFP;il34^+/-^* incrosses, or *rpe65a:nfsB-eGFP;csf1ra^+/-^*;*csf1rb^+/-^*incrosses were immersed in system water treated with 1.5X n-phenylthiourea (PTU; MilliporeSigma, St. Louis, MO, USA) from 0.25 – 5 dpf to prevent melanogenesis. On 5 dpf, larvae were screened for eGFP using an AxioZoom 16 Fluorescent Stereomicroscope (Zeiss, Oberkochen, Germany) and RPE was ablated by 24-hour immersion in 10 mM MTZ (MilliporeSigma) dissolved in system water. The MTZ solution was made on 5 dpf, immediately prior to screening, as described^30^. MTZ was washed out after 24 hours (designated 1- day post-injury (dpi)). Larvae euthanized ≤9 dpf remained in system water (unfed) until fixation, with daily water changes. Late-stage larvae euthanized on 19 dpf were washed an additional ≥3 times with system water after MTZ removal (e.g., between 6 – 7 dpf/1 – 2 dpi) and then put into rotifer polyculture on 7 dpf/2 dpi.

### *il34* plasmid generation and RNA probe synthesis

Primers were designed to encompass the full *Danio rerio il34* NCBI reference mRNA sequence length (NM_001128701.1) (Table S2). Fragment PCR was performed on a PTC-100 thermal cycler (Bio-Rad) using template cDNA generated from 4 dpi eGFP^+^ fluorescence-activated cell sorting (FACS)-isolated RPE. Alternatively, cDNA synthesized from enucleated 4 dpi eyes could be used. Full-length *il34* PCR fragments were gel-isolated and purified using the QIAquick Gel Extraction kit (Qiagen, Venlo, Netherlands). Fragments were cloned into the pGEM^®^-T Easy Vector System (Promega, Madison, WI, USA) and transformed into competent cells (DH5ɑ E. coli strain) according to manufacturer protocols. Sequencing was performed to identify correctly assembled pGEM-T-fl*il34* constructs. The confirmed pGEM-T-fl*il34* template was linearized using SacII (New England Biolabs) and purified using the QIAquick PCR Purification kit (Qiagen). Anti-sense RNA probes were synthesized by *in vitro* transcription using the DIG RNA Labeling Mix (Roche, Basel, Switzerland) protocol, with minimal modifications. Specifically, RNase OUT and dithiothreitol (DTT) were added to the 20 µL DIG RNA labeling mixture, which was incubated at 37 °C for 3 hours, followed by treatment with RNase-free DNaseI for 15 – 20 minutes at 37 °C. As described previously^73^, RNA probes were precipitated at-20 °C overnight in lithium chloride (LiCl_2_). The next day, precipitated RNA was pelleted by centrifugation (∼20,000 x *g* for 15 minutes at 4 °C), washed with ice cold 70% ethanol (RNase-free), pelleted again (∼20,000 x *g* for 15 minutes at 4 °C), and allowed to dry completely at room temperature. RNA probes were resuspended in RNase-free water and an equal volume of formamide was added for long-term storage at-20 °C.

### *il34 in situ* hybridization

Whole mount *in situ* hybridization was performed following previously described methods^73^, with modification. Briefly, 7 dpf zebrafish were euthanized and fixed in 4% paraformaldehyde (PFA) for 3 – 4 hours at room temperature or at 4 °C overnight. Post-fixation, larvae were dehydrated in 100% methanol and stored at-20 °C long-term. For experiments, 100% methanol-stored larval tissues were rehydrated through a gradient of 50% methanol in PSBT (1X PBS with 0.1% Tween-20) to 30% methanol in PSBT, followed by two PBST only washes. To depigment the RPE, larvae were bleached in a 3% hydrogen peroxide + 1% potassium hydroxide solution until RPE pigment was visibly diminished (using a stereomicroscope). Larvae were permeabilized using collagenase (1 mg/mL in PBST) for 15 minutes followed by Proteinase K (10 µg/mL in PBST) for 10 minutes, then rinsed twice with PBST. Larvae were refixed prior to hybridization in 4% PFA followed by two PBST washes. Prehybridization, and overnight probe hybridization was performed as described with the same hybridization buffer^73^, with one important difference – prior to adding to tissues, the diluted probe in hybridization buffer was incubated at 68 °C for 5 minutes. The next day, anti-DIG-AP Fab fragments antibody (Roche, RRID:AB_514497) was preabsorbed in blocking solution (1% bovine serum albumin, 1% dimethylsulfoxide (DMSO), 0.3% Triton X-100, 0.25% Tween 20, 2% normal goat serum (NGS) in 1X PBS) for at least 2 hours at 4 °C. Larvae were then taken through a series of washes: 50% formamide/2xSSC + 0.1% Tween 20 (diluted from pH 7 20X SSC (3M NaCL + 0.3M sodium citrate)) for 20 minutes at 55 °C, 3x 10 minutes at 37 °C in 2xSSC + 0.1% Tween 20 (SSCT), 2x 15 minutes at 55 °C in 0.2xSSCT, 5 minutes at 37 °C in PBST, and 5 minutes at room temperature in PBST. Larvae were blocked for at least 1 hour then incubated with preabsorbed anti-DIG antibody overnight at 4 °C. The following day, 4x 20-minute PBST and 3x 5-minute staining buffer (100mM Tris pH 9.5, 50mM MgCl_2_, 100mM NaCl, 0.1% Tween 20) washes were performed and NBT/BCIP (Roche) detection was done to desired staining level. When achieved, larvae were washed for 3x 10 minutes in PBST and preserved in 4% PFA at 4 °C. Post-labeling, larval eyes were either enucleated for wholemount imaging or larvae were washed out of fixation, sucrose-protected, embedded, and sectioned (12μm) as described below for immunostaining. Coverslips (No. 1) were mounted using DPX Mounting Medium (Electron Microscopy Sciences, Hatfield, PA, USA) and all *in situ* hybridization images were obtained with a 40X objective on a Leica DM2500 upright microscope (Wetzlar, Germany).

### TUNEL and BrdU incorporation assays

TUNEL assays were performed using the Click-iT Plus TUNEL Assay for In Situ Apoptosis Detection kit, Alexa Fluor^TM^ 647 dye (Invitrogen, Waltham, MA, USA) following the manufacturer experimental protocol for tissue sections, with one modification. Briefly, sectioned samples on slides were incubated with 0.5X Proteinase K solution to avoid over digestion. BrdU assays were performed by immersing larvae in a 10 mM BrdU (MilliporeSigma) solution (dissolved in system water) for 24 hours prior to fixation. BrdU assay antibody (Abcam, Waltham, MA, USA, RRID:AB_305426) information can be found in Table S2 and staining specifics can be found below in the methods for immunostaining.

### Larval tissue processing & immunostaining

Zebrafish larvae aged 6 – 9 dpf were euthanized and fixed in 4% PFA for 3 – 4 hours at room temperature or at 4 °C overnight. Late-stage 19 dpf larvae were fixed in 4% PFA at 4 °C overnight and, the following day, rotated at room temperature for 1 hour prior to PFA washout. Larvae were washed 3x 5 minutes in 1X PBS following fixation and then incubated in 1X PBS at 4 °C until genotypes were confirmed (<1 month). Tissue processing and staining for all larval ages was performed using previously described, established protocols (see SI Appendix^8^ for extensive methods). Briefly, genotyped larvae were sucrose-protected, embedded in optimal cutting temperature (OCT; Scigen, Singapore) compound, and cryosectioned at 12µm thickness onto poly-L-lysine-coated slides on a Leica CM1850 cryostat (Leica Biosystems, Wetzlar, Germany). Sections were rehydrated in 1X PBS for 5 minutes, blocked in 5% normal goat serum in PBTD (1X PBS with 0.1% Tween-20 and 1% DMSO) for ≥2 hours, and incubated in primary antibody at 4 °C overnight. For BrdU staining, rehydrated sections underwent antigen retrieval (4N HCl incubation for 8 minutes at 37 °C), then 3x 10-minute washes with PBTD prior to blocking. The next day, slides were washed 3x 10 minutes with PBTD and incubated in secondary antibody for 3 hours at room temperature. Slides were washed 3 additional times (10 minutes each) with PBTD, and nuclei were counterstained with DAPI. Coverslips (No. 1 or No. 1.5) were mounted using liquid Vectashield Antifade Mounting Medium with DAPI (Vector Laboratories, Burlingame, CA, USA) and sealed with nail polish. Slides were stored in the dark at 4 °C until imaging (≤1 week). All antibodies and dilutions used for this study can be found in Table S2.

### Confocal image acquisition & processing

Images were acquired using the following confocal microscopes: Olympus Fluoview FV1200 (Olympus Corporation, Tokyo, Japan), Nikon A1R and Nikon AXR (Nikon Corporation, Tokyo, Japan). Imaging on Nikon A1R and AXR microscopes was performed at the Center for Biomedical Research Support Microscopy and Imaging Facility at UT Austin (RRID:SCR_021756). Sectioned tissue on slides were imaged using 40X (1.30 NA) oil immersion lenses (Olympus or Nikon Corporations) and digital zooms (1.2–1.4X) were implemented for some experiments to focus on ocular tissues. All acquired z-stacks contained between 18–20 total slices with 1 μm z-step intervals. Raw imaging data were quantified and processed for presentation in figures using FIJI (ImageJ, RRID:SCR_002285)^74^ or Imaris for Neuroscientists (version 10.2; Bitplane, Belfast, United Kingdom, RRID:SCR_007370). 19 dpf/14 dpi eyes were usually large enough to require two images, which were stitched together for visualization, quantification, and data presentation purposes using the 3D Stitching plugin in FIJI^75^. For all imaging datasets, experimental replicates (N) indicate the number of times an experiment was performed, and biological replicates (n) represent the total number of individual larvae analyzed (collated in Table S1).

### Quantification of imaging data

#### Automated quantification (FIJI and MATLAB)

Automated quantification was preferentially implemented whenever possible to limit bias. Percent eGFP^+^ and mCherry^+^ area quantification was performed in FIJI on 8-bit maximum-projected z-stacks within RPE monolayer or whole eye regions of interest (ROIs) generated manually using the polygon selection tool, as previously described^8^. To further prevent bias, thresholding was set based on existing FIJI algorithms (e.g., Otsu, Triangle, Yen) that best fit the data for each individual animal/experiment. All percent area measurements were made from the central-most section of each larva. Automated quantification of TUNEL^+^ puncta was also performed in FIJI on 8-bit maximum-projected z-stacks within an ROI encompassing the ONL and RPE, and images were processed as previously described (SI Appendix^8^). The Analyze Particles tool was used to count puncta with the following parameters – size = 2 – infinity μm^2^; circularity = 0.00 – 1.00. Data from TUNEL experiments represent the sum of cells from three central cryosections, except for the *csf1ra/b* 6 dpf/1 dpi datasets, which, due to difficulty of getting double mutant tissue, represent the sum of cells from two central cryosections. Automated quantification of pigment recovery was performed in MATLAB (version R2022a; MathWorks, Natick, MA, USA, RRID:SCR_001622) using RpEGEN scripts^30^. Preprocessing of confocal microscope z-stack images for RpEGEN was performed in FIJI, as described^30^. Briefly, images were imported into FIJI, converted to maximum intensity projections, reoriented so that dorsal was up and distal was left, and saved as 8-bit TIFs. RPE regions of interest (ROIs) were also created in FIJI using the Polygon Selections tool, then RpEGEN.m and RpEGEN_PermPlot.m scripts were run in MATLAB. ROIs from RpEGEN.m output files were validated prior to running the RpEGEN_PermPlot.m script.

#### Manual quantification (FIJI)

4C4^+^ cells, mCherry^+^ cells, BrdU^+^ cells, and pyknotic nuclei were quantified manually using the FIJI Cell Counter plug-in, and data from each of these experiments represent the sum of cells from three central cryosections. Percent RPE regeneration was quantified as described previously (SI Appendix^8^), using the central-most section from each larva. Briefly, reference and midpoint lines were utilized to make dorsal and ventral angle measurements based on centrality of tissue regeneration. Angle measurements were then summed and converted to a percentage to determine the total amount of RPE regeneration (e.g. 100% = 180 angular degrees).

#### Imaris

The concept and strategy implemented to quantify depth of eGFP^+^ particle internalization was modeled from a previous study^76^. The 3 dpi *il34* dataset analyzed in Imaris represents a subset (N=1) of a larger (N=3) dataset used for eGFP percent area quantification (see Table S1). This dataset was chosen as it had three consecutive sections for each animal and minimal background and was, thus, most suitable for Imaris analyses. The mCherry detection channel, which labeled mononuclear phagocytes, was isosurfaced using the Surfaces automatic creation wizard. The eGFP channel was then masked to isolate eGFP^+^ signal that overlapped only with mCherry^+^ surfaces. The Surfaces automatic creation wizard was again implemented to isosurface the masked eGFP^+^ subchannel. For both channels, debris ≤10 µm^2^ was filtered out using the ‘filter surfaces’ function in the creation wizard. Particle internalization was then measured using the ‘Shortest Distance to Surfaces’ statistic, where Surfaces = mCherry (mononuclear phagocytes). Distance measurements were imported into Prism 10 (GraphPad Software, RRID:SCR_002798), where statistical tests were performed (Table S1). For percentages discussed in the Results section, particles with distance measurements ≤10 nm were determined to be at the surface, while particles with distance measurements ≥20 nm were determined to be engulfed.

#### Phenotype scoring

All 14 dpi *il34* and *csf1ra/b* late-stage larval phenotypes were scored blind using the following criteria. None = phenotype comparable to 19 dpf unablated controls, showing elongated photoreceptor outer segments (DAPI channel) that align parallel to and interdigitate with RPE apical microvilli (eGFP channel). Mild = presenting with minimal disorganization and/or truncation of outer segments and/or truncated RPE apical microvilli, no obvious pigment debris (brightfield channel). Moderate = presenting with highly disorganized and/or truncated outer segments and/or truncated apical microvilli, some smaller pigment debris that does not impede tissue regrowth. Severe = presenting with absence/loss of outer segment and/or RPE tissue, pigment debris that is visibly impeding tissue integrity.

### Statistics, data visualization, and exclusion criteria

Detailed statistical information for this study can be found in Table S1. Prism 10 (GraphPad Software) was used to perform statistical analyses, Pearson correlation analyses, and graphical representation of raw data. Normal (Gaussian) distribution in datasets was determined using the D’Agostino–Pearson omnibus normality test. For pairwise comparisons, significance values were calculated using unpaired t tests with Welch’s correction (normal distribution) or nonparametric Mann–Whitney U tests (nonnormal distribution). To determine variance and perform multiple significance comparisons, Welch’s ANOVA with Dunnett’s post hoc analysis (normal distribution) or nonparametric Kruskal–Wallis one-way ANOVA with Dunn’s post hoc analysis (nonnormal distribution) was performed. For all graphs, quartiles are delimited, and the median represents the measure of center (solid line). P-values ≤0.05 were considered significant and summarized as follows in all graphs: *, ≤0.05; **, ≤0.01; ***, ≤0.001; ****, ≤0.0001. In RpEGEN output graphs (Figure 1), 95% confidence intervals are represented by shaded envelopes (Figure 1A) and a dotted black line (Figure 1B). All color palettes used for graphical representation and figures were from (or inspired by) Paul Tol’s color schemes and templates^77^. Animals were excluded from analysis if tissue was of generally poor quality due to processing; folded, torn, or missing in critical areas; or if central sections could not be obtained for analysis. As mentioned above, 5 dpf eGFP^+^ siblings were randomly assigned to unablated or ablated experimental groups after screening.

## Supporting information

Supplemental Information compiled

## Acknowledgements

This work was supported by the BrightFocus Foundation New Investigator Grant Program in Macular Degeneration Research (M2023012N to L.L.L), the National Institutes of Health (RO1-EY29410 to J.M.G. and NIH Core Grant P30-EY08098 to the University of Pittsburgh Department of Ophthalmology), and Research to Prevent Blindness, Inc. Additional support was received from the Eye and Ear Foundation of Pittsburgh Wiegand Fellowship in Ophthalmology (to L.L.L). We thank Elli Davis for technical assistance; Rufus Fisher for editorial assistance; and the zebrafish teams at The University of Pittsburgh and UT Austin for exceptional facility maintenance, general support, and animal care.

## Author contributions

Conceptualization, L.L.L. and J.M.G.; Validation, L.L.L.; Formal Analysis, L.L.L.; Investigation, L.L.L., R.G.G., S.U.J., J.M.G.; Writing – Original Draft, L.L.L.; Writing – Review & Editing, L.L.L., R.G.G., S.U.J., and J.M.G.; Visualization, L.L.L.; Supervision, L.L.L. and J.M.G.; Project Administration, L.L.L.; Funding Acquisition; L.L.L. and J.M.G.

## Notes

### Competing Interest Statement

The authors have declared no competing interest.

