## Supplemental Information compiled for "Interleukin-34 and debris clearance by mononuclear phagocytes drive retinal pigment epithelium regeneration in zebrafish"

**Table S1.** Experimental groups and statistical information for this study.

| *Gene* | **Experimental groups** | **# experiments (N) / biological replicates (n)** | **Experimental analysis** | **Statistical test (post hoc test)** | ***p-value*** | ***p-value* summary** | **Figure shown** |
|---|---|---|---|---|---|---|---|
| *il34* | | | | | | | |
| * | 9dpf +/+ & 9dpf -/- (MTZ-) | **5** / n=18 (+/+) & n=17 (-/-) | RpEGEN (MATLAB) | None | – | – | Figure 1A Figure S2C,D |
| * | 4dpi +/+ vs 4dpi -/- (MTZ+) | **5** / n=22 (+/+) vs n=22 (-/-) | RpEGEN (MATLAB) | RpEGEN_PermPlot.m | various | various | Figure 1A,B |
| * | 4dpi +/+ vs 4dpi -/- (MTZ+) | **5** / n=22 (+/+) vs n=22 (-/-) | eGFP (FIJI) | Welch's ANOVA (Dunnett's) | 0.4754 | ns | Figure 2H |
| | 6dpf +/+ vs 6dpf -/- (MTZ-) | **3** / n=17 (+/+) vs n=17 (-/-) | TUNEL (FIJI) | Kruskal–Wallis (Dunn's) | >0.9999 | ns | Figure S3C Figure S2A,B |
| | 1dpi +/+ vs 1dpi -/- (MTZ+) | **3** / n=19 (+/+) vs n=17 (-/-) | TUNEL (FIJI) | Kruskal–Wallis (Dunn's) | >0.9999 | ns | Figure S3A-C |
| † | 7dpf +/+ vs 7dpf -/- (MTZ-) | **2** / n=10 (+/+) vs n=8 (-/-) | 4C4 (manual) | Welch's t-test | 0.0866 | ns | Figure S4A-C |
| † | 7dpf +/+ vs 7dpf -/- (MTZ-) | **3** / n=15 (+/+) vs n=13 (-/-) | mCherry (manual) | Welch's t-test | 0.1043 | ns | Figure S4C |
| | 8dpf +/+ vs 8dpf -/- (MTZ-) | **3** / n=12 (+/+) vs n=10 (-/-) | mCherry (FIJI) | Welch's t-test | 0.1696 | ns | Figure S4D-F |
| ‡ | 2dpi +/+ vs 2dpi -/- (MTZ+) | **2** / n=13 (+/+) vs n=13 (-/-) | 4C4 (manual) | Welch's t-test | <0.0001 | **** | Figure 2A-C |
| ‡ | 2dpi +/+ vs 2dpi -/- (MTZ+) | **3** / n=14 (+/+) vs n=11 (-/-) | mCherry (manual) | Welch's t-test | 0.0018 | ** | Figure 2D |
| | 3dpi +/+ vs 3dpi -/- (MTZ+) | **3** / n=17 (+/+) vs n=27 (-/-) | mCherry (FIJI) | Welch's t-test | 0.2249 | ns | Figure 2E-G |
| ‡ | 2dpi +/+ vs 2dpi -/- (MTZ+) | **5** / n=27 (+/+) vs n=24 (-/-) | eGFP (FIJI) | Welch's ANOVA (Dunnett's) | 0.4092 | ns | Figure 2A,B,H |
| § | 3dpi +/+ vs 3dpi -/- (MTZ+) | **3** / n=17 (+/+) vs n=27 (-/-) | eGFP (FIJI) | Welch's ANOVA (Dunnett's) | 0.0009 | *** | Figure 2E,F,H |
| § | 3dpi +/+ vs 3dpi -/- (MTZ+) | n=720 (+/+) vs n=870 (-/-) (subset of above) | shortest distance (Imaris) | Welch's t-test | <0.0001 | **** | Figure S5A |
| | 2dpi +/+ vs 3dpi +/+ (MTZ+) | **see above** / n=27 (2dpi) vs n=17 (3dpi) | eGFP (FIJI) | Welch's ANOVA (Dunnett's) | <0.0001 | **** | Figure 2H |
| | 3dpi +/+ vs 4dpi +/+ (MTZ+) | **see above** / n=17 (3dpi) vs n=22 (4dpi) | eGFP (FIJI) | Welch's ANOVA (Dunnett's) | 0.0019 | ** | Figure 2H |
| | 2dpi -/- vs 3dpi -/- (MTZ+) | **see above** / n=24 (2dpi) vs n=27 (3dpi) | eGFP (FIJI) | Welch's ANOVA (Dunnett's) | 0.6176 | ns | Figure 2H |
| | 3dpi -/- vs 4dpi -/- (MTZ+) | **see above** / n=27 (3dpi) vs n=22 (4dpi) | eGFP (FIJI) | Welch's ANOVA (Dunnett's) | <0.0001 | **** | Figure 2H |
| † | 7dpf +/+ vs 7dpf -/- (MTZ+) | **5** / n=25 (+/+) vs n=21 (-/-) | pyknotic nuclei (manual) | Welch's ANOVA (Dunnett's) | 0.0208 | * | Figure 3C |
| ‡ | 2dpi +/+ vs 2dpi -/- (MTZ+) | **5** / n=27 (+/+) vs n=24 (-/-) | pyknotic nuclei (manual) | Welch's ANOVA (Dunnett's) | <0.0001 | **** | Figure 3A-C |
| | 8dpf +/+ vs 8dpf -/- (MTZ-) | **3** / n=17 (+/+) vs n=19 (-/-) | TUNEL (FIJI) | Welch's ANOVA (Dunnett's) | 0.9658 | ns | Figure 3F |
| | 3dpi +/+ vs 3dpi -/- (MTZ+) | **3** / n=22 (+/+) vs n=21 (-/-) | TUNEL (FIJI) | Welch's ANOVA (Dunnett's) | <0.0001 | **** | Figure 3D-F |
| * | 9dpf +/+ vs 9dpf -/- (MTZ-) | **2** / n=9 (+/+) vs n=16 (-/-) | pyknotic nuclei (manual) | Welch's ANOVA (Dunnett's) | 0.9996 | ns | Figure 3I |
| * | 4dpi +/+ vs 4dpi -/- (MTZ+) | **2** / n=12 (+/+) vs n=14 (-/-) | pyknotic nuclei (manual) | Welch's ANOVA (Dunnett's) | >0.9999 | ns | Figure 3G-I |

| | | | | | | | |
|---|---|---|---|---|---|---|---|
| | 8dpf +/+ vs 8dpf -/- (MTZ-) | **3** / n=21 (+/+) vs n=18 (-/-) | BrdU (manual) | Welch's ANOVA (Dunnett's) | >0.9999 | ns | Figure S6E |
| | 3dpi +/+ vs 3dpi -/- (MTZ+) | **3** / n=13 (+/+) vs n=21 (-/-) | BrdU (manual) | Welch's ANOVA (Dunnett's) | 0.0585 | ns | Figure S6A,B,E |
| | 9dpf +/+ vs 9dpf -/- (MTZ-) | **5** / n=23 (+/+) vs n=18 (-/-) | BrdU (manual) | Welch's ANOVA (Dunnett's) | 0.9999 | ns | Figure S6F |
| | 4dpi +/+ vs 4dpi -/- (MTZ+) | **5** / n=19 (+/+) vs n=23 (-/-) | BrdU (manual) | Welch's ANOVA (Dunnett's) | >0.9999 | ns | Figure S6C,D,F |
| | 19dpf +/+ & 19dpf -/- (MTZ-) | **1** / n=6 (+/+) & n=6 (-/-) | visualization / phenotype scoring | None | – | – | Figure 4B |
| | 14dpi +/+ & 14dpi -/- (MTZ+) | **1** / n=6 (+/+) & n=6 (-/-) | visualization / phenotype scoring | None | – | – | Figure 4C-E |
| *csfr1a/ csf1rb* | 6dpf +/+ vs 6dpf DM (MTZ-) | **4** / n=14 (+/+) vs n=12 (DM) | TUNEL (FIJI) | Welch's ANOVA (Dunnett's) | 0.3790 | ns | Figure S3F Figure S2E,F |
| | 1dpi +/+ vs 1dpi DM (MTZ+) | **3** / n=9 (+/+) vs n=12 (DM) | TUNEL (FIJI) | Welch's ANOVA (Dunnett's) | 0.2137 | ns | Figure S3D-F |
| [¶] | 8dpf +/+ vs 8dpf DM (MTZ-) | **2** / n=8 (+/+) vs n=6 (DM) | 4C4 (manual) | Mann-Whitney test | 0.0003 | *** | Figure S4G-I |
| [#, Δ] | 3dpi +/+ vs 3dpi DM (MTZ+) | **3** / n=13 (+/+) vs n=18 (DM) | 4C4 (manual) | Welch's t-test | <0.0001 | **** | Figure 5A-C |
| [#, Δ] | 3dpi +/+ vs 3dpi DM (MTZ+) | **5** / n=31 (+/+) vs n=29 (DM) | eGFP (FIJI) | Mann-Whitney test | <0.0001 | **** | Figure 5A,B,D |
| [¶] | 8dpf +/+ vs 8dpf DM (MTZ-) | **4** / n=22 (+/+) vs n=19 (DM) | pyknotic nuclei (manual) | Kruskal–Wallis (Dunn's) | 0.7662 | ns | Figure 5E |
| [#, Δ] | 3dpi +/+ vs 3dpi DM (MTZ+) | **4** / n=26 (+/+) vs n=24 (DM) | pyknotic nuclei (manual) | Kruskal–Wallis (Dunn's) | 0.0001 | *** | Figure 5A,B,E |
| | 9dpf +/+ & 9dpf DM (MTZ-) | **2** / n=4 (+/+) & n=7 (DM) | visualization | None | – | – | Figure S2G,H |
| | 4dpi +/+ vs 4dpi DM (MTZ+) | **2** / n=11 (+/+) vs n=11 (DM) | % regeneration (manual) | Welch's t-test | <0.0001 | **** | Figure 5F-H |
| | 19dpf +/+ & 19dpf DM (MTZ-) | **2** / n=6 (+/+) & n=5 (DM) | visualization / phenotype scoring | None | – | – | Figure 6C |
| | 14dpi +/+ & 14dpi DM (MTZ+) | **3** / n=12 (+/+) & n=6 (DM) | visualization / phenotype scoring | None | – | – | Figure 6A,D-G |
| wildtype | 7dpf (MTZ-) | **1** / n=10 | *in situ* hybridization *(il34)* | None | – | – | Figure S1A,C |
| | 2dpi (MTZ+) | **1** / n=8 | *in situ* hybridization *(il34)* | None | – | – | Figure S1B,D |

Abbreviations: DM, double mutant; dpf, days post-fertilization; dpi, days post-injury; FIJI, FIJI is just ImageJ; MTZ, metronidazole; ns, not significant.

[*] 4dpi *il34* datasets used for RpEGEN were also used for eGFP % area quantification. One 9dpf/4dpi RpEGEN dataset with 3 central sections/animal was used for pyknotic counts.

[†] 7dpf *il34* 4C4 and mCherry datasets were pooled and used for pyknotic nuclei counts.

[‡] 2dpi *il34* 4C4 and mCherry datasets were pooled and used for pyknotic nuclei counts and eGFP % area quantification.

[§] 3dpi *il34* Imaris dataset represents a subset (N=1) of the larger eGFP % area quantification dataset. The dataset had three consecutive sections for each animal and minimal background and was, thus, most suitable for Imaris analyses.

[¶] The 8dpf *csf1ra/csf1rb* 4C4 dataset was also used for pyknotic nuclei counts.

[#] Four of the 3dpi *csf1ra/csf1rb* eGFP % area quantification datasets were also used for pyknotic nuclei counts. Three were also used for 4C4 cell counts.

[Δ] N=2 experiments overlapping for 4C4, eGFP, and pyknotic nuclei counts were used for the Pearson correlation test (n=9 +/+ & n=13 DM). Pearson's correlation r and p-value results are in Figure S7.

**Table S2.** Resources and reagents used for this study.

| Reagent type | Name/Identifier | Manufacturer/Resource info | Catalog # | Other info (Reference) |
|---|---|---|---|---|
| Antibody (primary) | 4C4 (mouse monoclonal) | A kind gift from Dr. Peter Hitchcock, University of Michigan, Ann Arbor, MI | N/A | 1:200 (1) |
| Antibody (primary) | anti-mCherry (mouse monoclonal) | Takara Bio USA Inc./Clontech Laboratories | 632543 RRID:AB_2307319 | 1:200 |
| Antibody (primary) | anti-BrdU (rat monoclonal) | Abcam | ab6326 RRID:AB_305426 | 1:250 |
| Antibody (primary) | zpr-1 (mouse monoclonal) | Zebrafish International Resource Center (ZIRC) | ZDB-ATB-081002-43 | 1:250 |
| Antibody (secondary) | goat anti-mouse IgG Cy3 | Jackson ImmunoResearch Labs, West Grove, Pennsylvania | 115-165-166 RRID:AB_2338692 | 1:250 |
| Antibody (secondary) | goat anti-mouse IgG Cy5 | Jackson ImmunoResearch Labs | 115-175-166 RRID:AB_2338714 | 1:250 |
| Antibody (secondary) | goat anti-rat IgG Cy3 | Jackson ImmunoResearch Labs | 112-165-003 RRID:AB_2338240 | 1:250 |
| Antibody (*in situ*) | Anti-Digoxigenin-AP, Fab fragments | Millipore Sigma (Roche) Anti-Digoxigenin-AP, Fab fragments | 11093274910 RRID:AB_514497 | |
| Assay/kit | DIG RNA Labeling Mix | Millipore Sigma (Roche) | 11277073910 | |
| Assay/kit | Click-iT Plus TUNEL Assay for In Situ Apoptosis Detection | Invitrogen | C10619 | 647 dye kit |
| Chemical | Metronidazole (MTZ) | Millipore Sigma | M3761 | 10 mM working |
| Chemical | n-phenylthiourea (PTU) | Millipore Sigma | P7629 | 25X stock 1.5X working (2) |
| Chemical | Paraformaldehyde (16% w/v) methanol free | Fisher Scientific | AA433689M | 4% working |
| Chemical | 2-(4-Amidinophenyl)-6-indolecarbamidine dihydrochloride (DAPI) | Millipore Sigma | D9542 | 1 mg/mL stock 1:250 working |
| Chemical | VECTASHIELD Antifade Mounting Medium with DAPI | Vector Laboratories | H-1200 | |
| Chemical | Tricaine-S (MS-222) | Pentair Aquatic Ecosystems, Inc. | TRS1 | |
| Chemical | Dithiothreitol (DTT) | ThermoFisher Scientific | 70-726-5ML | 0.1 M stock |
| Chemical | NBT/BCIP | Millipore Sigma (Roche) | 11681451001 | |
| Chemical | DPX Mountant | Electron Microscopy Sciences | 13510 | |
| Chemical | 5-Bromo-2'-deoxyuridine (BrdU) | Millipore Sigma | B5002 | 10 mM working |
| Enzyme | HpyCH4IV | New England Biolabs | R0619 | |
| Enzyme | SpeI-HF | New England Biolabs | R3133 | |
| Enzyme | MspI | New England Biolabs | R0106 | |
| Enzyme | SacII | New England Biolabs | R0157 | |
| Enzyme | DNase I recombinant, RNase-free | Millipore Sigma (Roche) | 4716728001 | |
| Enzyme | RNaseOUT | ThermoFisher Scientific | 10777019 | |
| Enzyme | Proteinase K Solution, RNA grade | Invitrogen | 25530049 | 20 mg/mL stock 10 μg/mL working |
| Enzyme | Collagenase from *Clostridium histolyticum* | Millipore Sigma | C9891 | 1 mg/mL working |
| Primers (genotyping) | *re03 (il34)* forward *re03 (il34)* reverse | CAGGGCATTAAGAAGGTCTTAC CAAATGATATCATTGTTCTAAC | custom | HpyCH4IV cuts mutant |

| | | | | |
|---|---|---|---|---|
| Primers (genotyping) | *j4e1 (csf1ra)* forward<br><br>*j4e1 (csf1ra)* reverse | TCTGGGCAAAGAGGACAACATCAC-ACTA<br>CAAACCTTGCAGAGCTGTGG | custom | SpeI cuts wildtype |
| Primers (genotyping) | *re01 (csf1rb)* forward<br>*re01 (csf1rb)* reverse | GGACAGAGTTTTCGCTCCAG<br>ATTGGACTCCGCTCATGTTC | custom | MspI cuts wildtype |
| Primers *(in situ)* | full length *il34* forward<br>full length *il34* reverse | GGACGCGCGGAGAGAGCT<br>GGTAGTGAGAAGTTTATTCCCTAA | custom | |
| Recombinant DNA | pGEM-T-Easy vector | Promega | A1360 | |
| Serum | Goat serum | Millipore Sigma | G9023 | 5% working |
| Software/ Scripts | MATLAB | MathWorks<br>Toolboxes needed to run RpEGEN: Image Processing Toolbox, Curve Fitting Toolbox, Statistics and Machine Learning Toolbox | https://www.mathworks.com/products/get-matlab.html | |
| Software/ Scripts | RpEGEN | Created by Dr. G. Burch Fisher, University of Maryland Center for Environmental Science, Frostburg, MD; available for download at github.com/burchfisher/RpEGEN | N/A | (2) |
| Zebrafish line | *re03* | Generated by Dr. Tjakko van Ham, Erasmus University Medical Center Rotterdam, Netherlands; received as a kind gift from Dr. Celia Shiau, University of North Carolina at Chapel Hill | RRID:ZFIN_ZDB-GENO-200116-7 | (3) |
| Zebrafish line | *j4e1* | Generated by Dr. Steven Johnson, Washington University Medical School, St. Louis, MO; received as a kind gift from Dr. Florence Marlow, Icahn School of Medicine at Mount Sinai, New York, NY | RRID:ZFIN_ZDB-GENO-001205-7 | (4) |
| Zebrafish line | *re01* | Generated by Dr. Tjakko van Ham, Erasmus University Medical Center Rotterdam, Netherlands; received as a kind gift from Dr. Florence Marlow, Icahn School of Medicine at Mount Sinai, New York, NY | RRID:ZFIN_ZDB-GENO-190430-1 | (5) |
| Zebrafish line | *rpe65a:*nfsB-eGFP (mw86Tg) | A kind gift from Drs. Brian Link and Ross Collery, Medical College of Wisconsin, Milwaukee, WI | N/A | (6) |
| Zebrafish line | *mpeg1:*mCherry (gl23Tg) | Available at Zebrafish International Resource Center (ZIRC); received as a kind gift from Dr. Neil Hukriede, University of Pittsburgh, Pittsburgh, PA | ZL9939<br>RRID:ZIRC_ZL9939 | (7) |
| Zebrafish line | AB (wildtype) | Zebrafish International Resource Center (ZIRC) | ZL1<br>RRID:ZIRC_ZL1 | |

#### Supplemental Figure Legends

**Figure S1. *il34* expression is upregulated in the RPE injury site.** Micrographs showing RPE-specific *il34* expression in **(A,C)** 7 dpf unablated larvae ($MTZ^-$; *n=5* sectioned tissue, *n=5* wholemount tissue) and **(B,D)** 2 dpi ablated larvae ($MTZ^+$; *n=5* sectioned tissue, *n=3* wholemount tissue). *Black arrows* point to the RPE layer. Scale bars = 40µm. Abbreviations = dpf, days post-fertilization; dpi, days post-injury; MTZ, metronidazole; RPE, retinal pigment epithelium. Additional experimental information can be found in Table S1.

**Figure S2. The RPE develops normally in unablated *il34*$^{-/-}$ and *csf1r*$^{DM}$ zebrafish. (A–D)** Transverse confocal micrographs showing endogenous *rpe65a:nfsB-eGFP* transgene expression *(green)* in unablated ($MTZ^-$) **(A)** 6 dpf *il34*$^{+/+}$, **(B)** 6 dpf *il34*$^{-/-}$, **(C)** 9 dpf *il34*$^{+/+}$, and **(D)** 9 dpf *il34*$^{-/-}$ larvae. **(E–H)** Transverse confocal micrographs showing endogenous *rpe65a:nfsB-eGFP* transgene expression *(green)* in unablated ($MTZ^-$) **(E)** 6 dpf *csf1r*$^{+/+}$, **(F)** 6 dpf *csf1r*$^{DM}$, **(G)** 9 dpf *csf1r*$^{+/+}$, and **(E)** 9 dpf *csf1r*$^{DM}$ larvae. **(a'–h''')** Digital zooms highlight the central RPE from three siblings per genotype and age group. **(a')** *White arrowheads* point to examples of normal RPE apical microvilli (present in all animals). DAPI *(white)* labels nuclei. Scale bars = 40µm; digital zoom scale bars = 20µm. Abbreviations = DM, double mutant; dpf, days post-fertilization; dpi, days post-injury; MTZ, metronidazole; RPE, retinal pigment epithelium. Experimental replicate information can be found in Table S1.

**Figure S3. *il34*$^{-/-}$ and *csf1r*$^{DM}$ zebrafish do not show more severe initial RPE ablation phenotypes.** Transverse confocal micrographs showing TUNEL labeling *(white)* in ablated ($MTZ^+$) **(A)** *il34*$^{+/+}$, **(B)** *il34*$^{-/-}$, **(D)** *csf1r*$^{+/+}$, and **(E)** *csf1r* $^{DM}$ larvae at 1 dpi. *Green lines* designate regions of interest (ROI) encompassing the photoreceptors and RPE. **(C)** Violin plots showing no significant differences between unablated (6 dpf, $MTZ^-$) or ablated (1 dpi, $MTZ^+$) *il34*$^{+/+}$ and *il34*$^{-/-}$ larvae ($p>0.9999$ and $p>0.9999$, respectively). **(F)** Violin plots showing no significant differences between unablated (6 dpf, $MTZ^-$) or ablated (1 dpi, $MTZ^+$) *csf1r*$^{+/+}$ and *csf1r*$^{DM}$ larvae ($p=0.3790$ and $p=0.2137$, respectively). Scale bars = 40µm. Abbreviations = DM, double mutant; dpf, days post-fertilization; dpi, days post-injury; MTZ, metronidazole; ns, not significant; ONL, outer nuclear layer; RPE, retinal pigment epithelium. Experimental replicates and statistical information can be found in Table S1.

**Figure S4. Ocular mononuclear phagocytes are present in unablated *il34*$^{-/-}$ and *csf1r*$^{DM}$ zebrafish. (A,B)** Transverse confocal micrographs showing 4C4 *(magenta)* labeling in unablated ($MTZ^-$) **(A)** *il34*$^{+/+}$ and **(B)** *il34*$^{-/-}$ larvae at 7 dpf. **(C)** Violin plots showing cell counts using two mononuclear phagocyte (MNP) markers – 4C4 (ns, $p=0.0866$) and *mpeg1:*mCherry (ns, $p=0.1034$) – in unablated ($MTZ^-$) *il34*$^{-/-}$ compared to *il34*$^{+/+}$ larvae at 7 dpf. **(D,E)** Transverse confocal micrographs showing *mpeg1:*mCherry *(magenta)* labeling in unablated ($MTZ^-$) **(D)** *il34*$^{+/+}$ and **(E)** *il34*$^{-/-}$ larvae at 8 dpf. **(F)** Violin plots showing no significant difference in mCherry expression (percent area) in unablated ($MTZ^-$) *il34*$^{+/+}$ and *il34*$^{-/-}$ at 8 dpf ($p=0.1696$). **(G,H)** Transverse confocal micrographs showing 4C4 *(magenta)* labeling in unablated ($MTZ^-$) **(G)** *csf1r*$^{+/+}$ and **(H)** *csf1r*$^{DM}$ larvae at 8 dpf. **(I)** Violin plots showing significantly less ocular 4C4-positive cells in unablated ($MTZ^-$) *csf1r*$^{DM}$ larvae at 8 dpf ($p=0.0003$). Digital zoom insets *(cyan outlines)* further emphasize MNPs ($4C4^+$ or $mCherry^+$ cells) without eGFP overlay. DAPI *(white)* labels nuclei. Scale bars = 40µm; *inset* scale bars = 20µm. Abbreviations = DM, double mutant; dpf, days post-fertilization; dpi, days post-injury; MTZ, metronidazole; ns, not significant; RPE, retinal pigment epithelium. Experimental replicates and statistical information can be found in Table S1.

**Figure S5. Mononuclear phagocytes in *il34*$^{+/+}$ and *il34*$^{-/-}$ larvae internalize eGFP$^{+}$ particles. (A)** Scatter plots showing significantly less internalization of eGFP$^{+}$ particles in 3 dpi ablated (MTZ$^{+}$) *il34*$^{-/-}$ larvae compared to *il34*$^{+/+}$ controls. *Black dotted lines* represent the mean. *Solid gray line* at y=0.0 is analogous to the MNP surface, depicted by the cartoon (created in https://BioRender.com). **(B)** 3D rendering of an ablated (MTZ$^{+}$) *il34*$^{+/+}$ mCherry$^{+}$ MNP *(magenta)* with internalized particles. *Color bar* indicates shortest distance of particle from the MNP surface. Scale bar represents 5μm; ****, p≤0.0001. Experimental replicates and statistical information can be found in Table S1.

**Figure S6. *il34*$^{-/-}$ zebrafish do not show proliferation phenotypes post-ablation. (A–D)** Transverse confocal micrographs showing BrdU labeling *(white)* in ablated (MTZ$^{+}$) *il34*$^{+/+}$ and *il34*$^{-/-}$ larvae at **(A,B)** 3 dpi and **(C,D)** 4 dpi. Digital zooms *(yellow outlines)* and *arrowheads* highlight BrdU$^{+}$ cells in the RPE layer. *Solid cyan lines* outline the lens and outer limits (sclera) of the eye. **(E)** Violin plots showing no significant difference between unablated (8 dpf, MTZ$^{-}$) or ablated (3 dpi, MTZ$^{+}$) *il34*$^{+/+}$ and *il34*$^{-/-}$ larvae ($p$>0.9999 and $p$=0.0585, respectively). **(F)** Violin plots showing no significant difference between unablated (9 dpf, MTZ$^{-}$) or ablated (4 dpi, MTZ$^{+}$) *il34*$^{+/+}$ and *il34*$^{-/-}$ larvae ($p$=0.9999 and $p$>0.9999, respectively). Scale bars = 40μm; *inset* scale bars = 20μm. Abbreviations = dpf, days post-fertilization; dpi, days post-injury; MTZ, metronidazole; ns, not significant; RPE, retinal pigment epithelium. Experimental replicates and statistical information can be found in Table S1.

**Figure S7. *csf1ra/b* 3 dpi RPE show very strong and highly significant negative correlations between microglia density and debris accumulation.** Pearson correlation matrix showing the relationships between 4C4, eGFP$^{+}$ debris, and pyknotic nuclei in a collective *csf1r*$^{+/+}$ and *csf1r*$^{DM}$ 3 dpi dataset. *Color bar* and *black text* in the matrix correspond to the Pearson correlation coefficient (r). *White text* indicates p-values. $r^2$ values are as follows: 4C4, eGFP ($r^2$ = 0.64); 4C4, pyknotic nuclei ($r^2$ = 0.6889); eGFP, pyknotic nuclei ($r^2$ = 0.6084). Experimental replicate information can be found in Table S1.

**Figure S1**

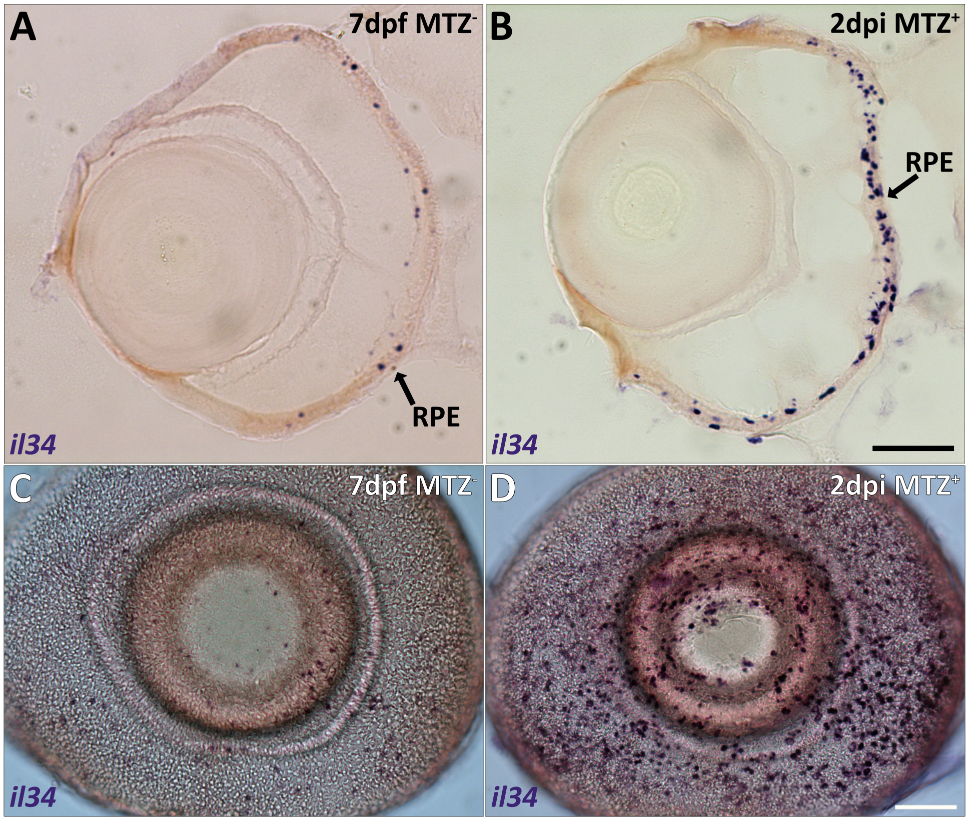

A
7dpf MTZ⁻
RPE
il34
B
2dpi MTZ⁺
RPE
il34
C
7dpf MTZ⁻
il34
D
2dpi MTZ⁺
il34

**Figure S2**

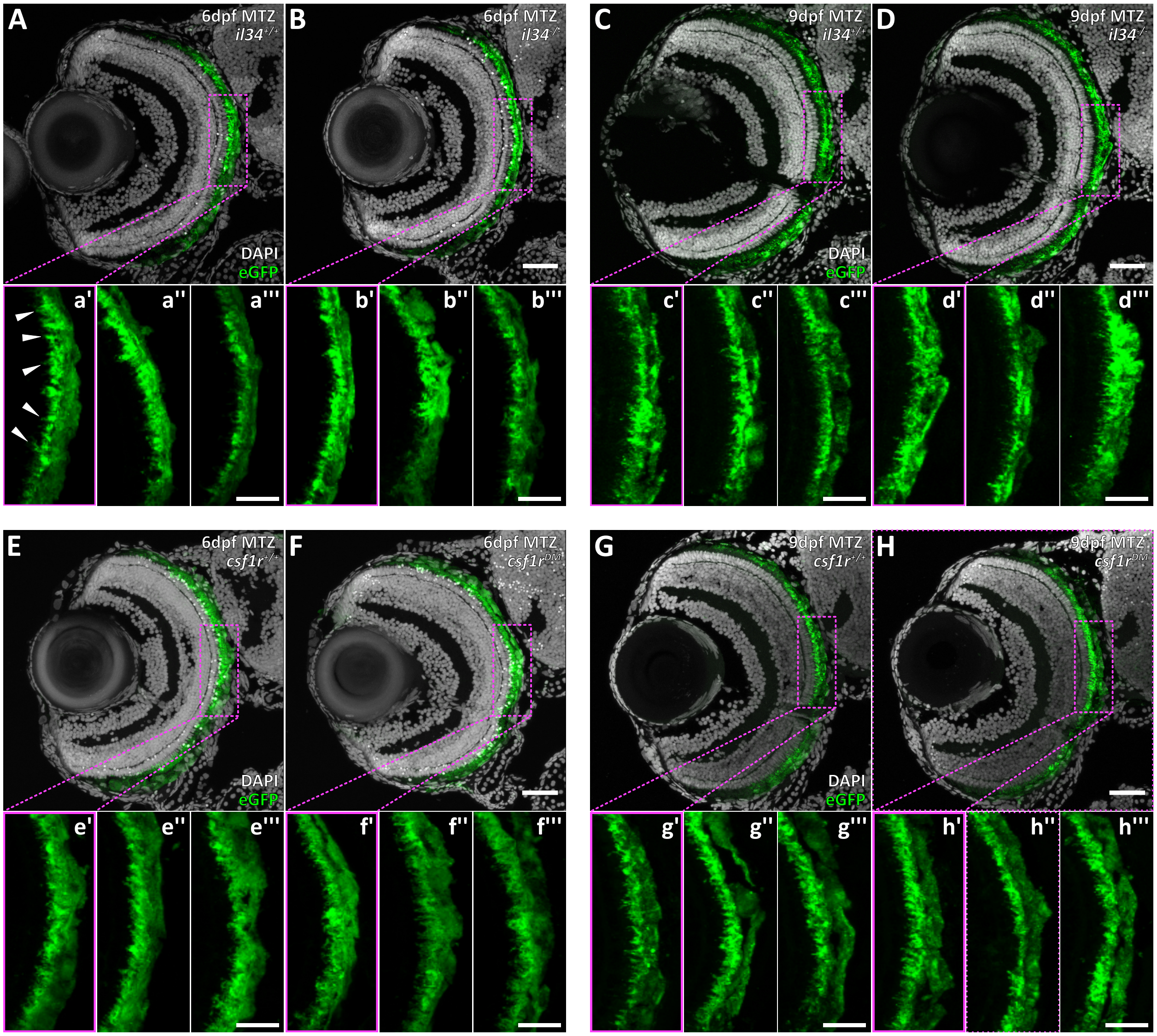

A
6dpf MTZ
$il34^{+/+}$
DAPI
eGFP
B
6dpf MTZ
$il34^{-/-}$
C
9dpf MTZ
$il34^{+/+}$
DAPI
eGFP
D
9dpf MTZ
$il34^{-/-}$
a'
a''
a'''
b'
b''
b'''
c'
c''
c'''
d'
d''
d'''
E
6dpf MTZ
$csf1r^{+/+}$
DAPI
eGFP
F
6dpf MTZ
$csf1r^{DM}$
G
9dpf MTZ
$csf1r^{+/+}$
DAPI
eGFP
H
9dpf MTZ
$csf1r^{DM}$
e'
e''
e'''
f'
f''
f'''
g'
g''
g'''
h'
h''
h'''

### Figure S3

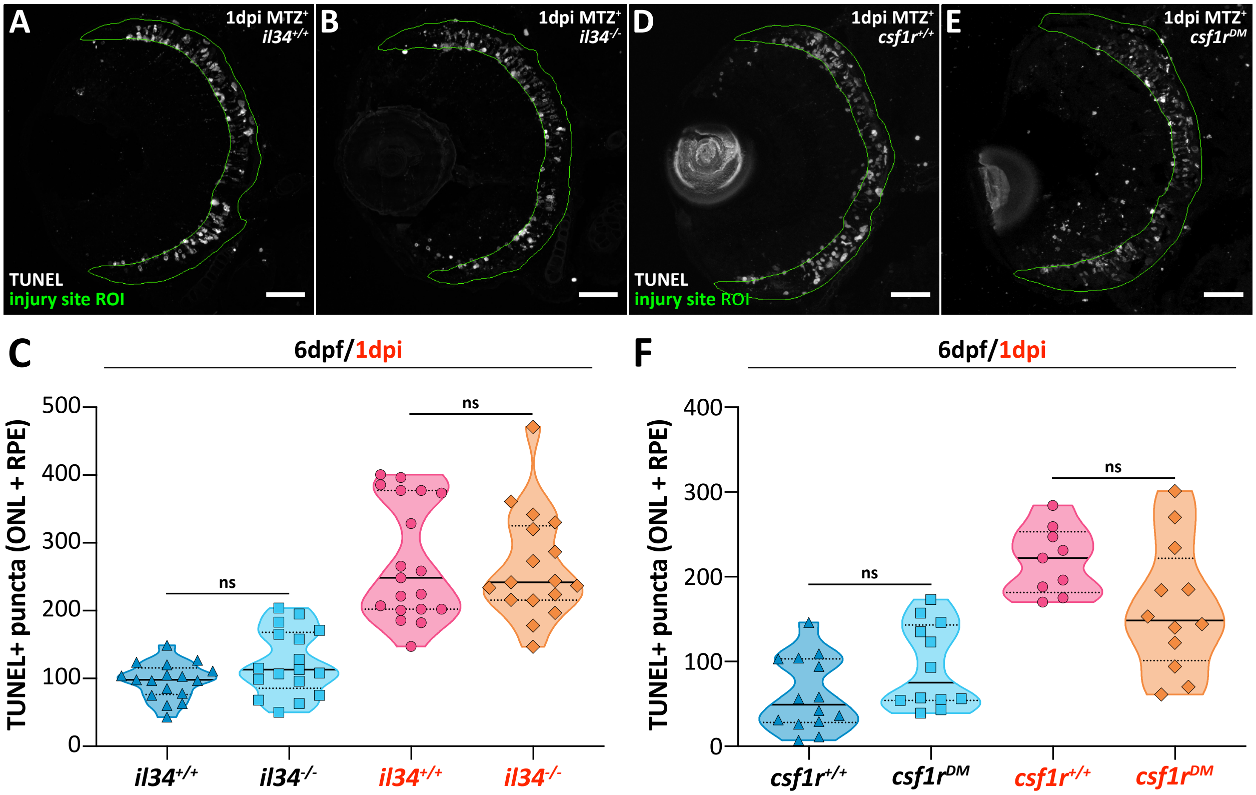
A
1dpi MTZ+ il34+/+
TUNEL
injury site ROI
B
1dpi MTZ+ il34-/-
D
1dpi MTZ+ csf1r+/+
TUNEL
injury site ROI
E
1dpi MTZ+ csf1rDM
C
6dpf/1dpi
TUNEL+ puncta (ONL + RPE)
500
400
300
200
100
0
ns
ns
il34+/+
il34-/-
il34+/+
il34-/-
F
6dpf/1dpi
TUNEL+ puncta (ONL + RPE)
400
300
200
100
0
ns
ns
csf1r+/+
csf1rDM
csf1r+/+
csf1rDM

### Figure S4

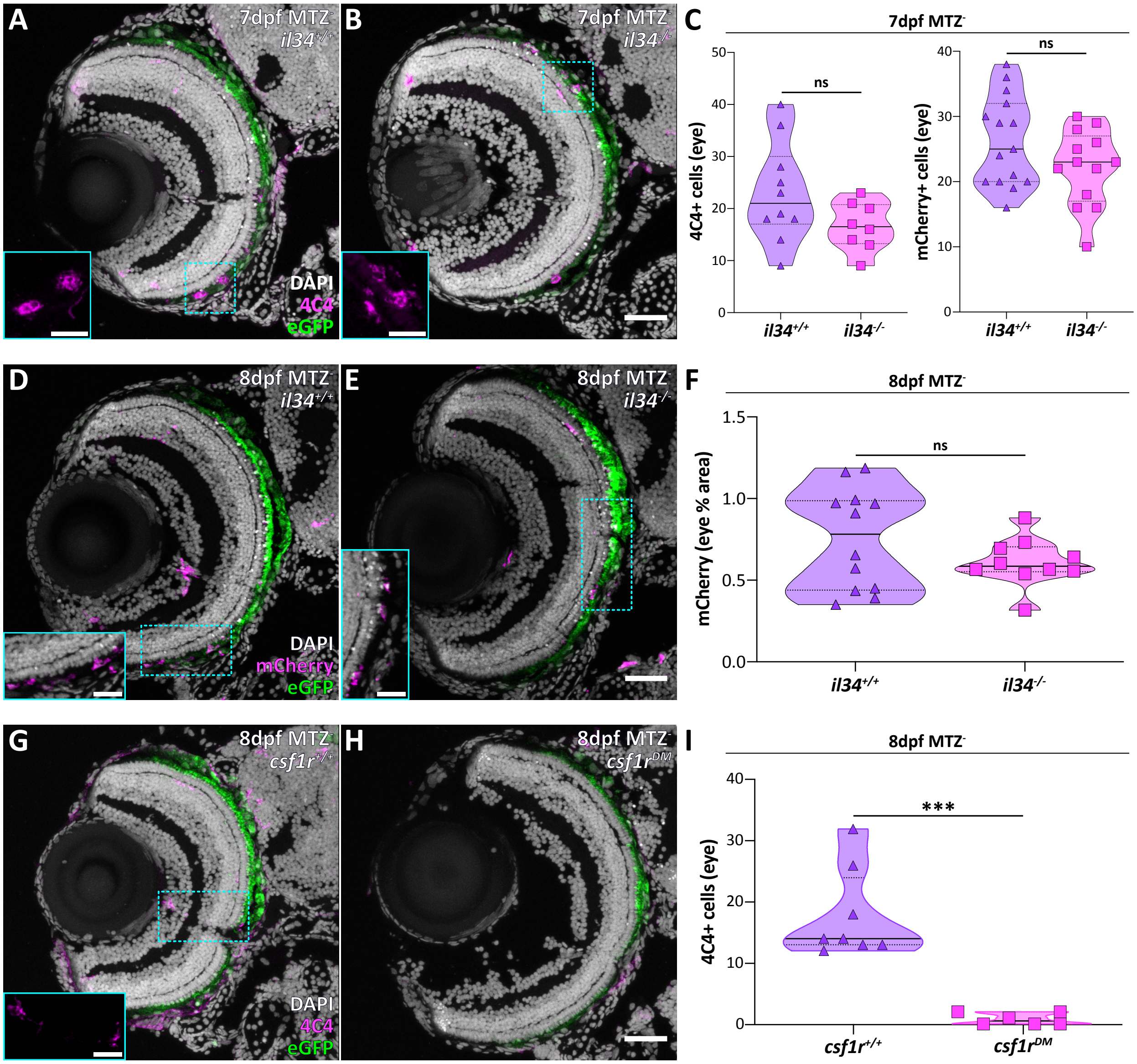

A
7dpf MTZ-
il34+/+
DAPI
4C4
eGFP
B
7dpf MTZ-
il34-/-
C
7dpf MTZ-
4C4+ cells (eye)
mCherry+ cells (eye)
ns
ns
il34+/+
il34-/-
D
8dpf MTZ-
il34+/+
DAPI
mCherry
eGFP
E
8dpf MTZ-
il34-/-
F
8dpf MTZ-
mCherry (eye % area)
ns
il34+/+
il34-/-
G
8dpf MTZ-
csf1r+/+
DAPI
4C4
eGFP
H
8dpf MTZ-
csf1rDM
I
8dpf MTZ-
4C4+ cells (eye)
***
csf1r+/+
csf1rDM

### Figure S5

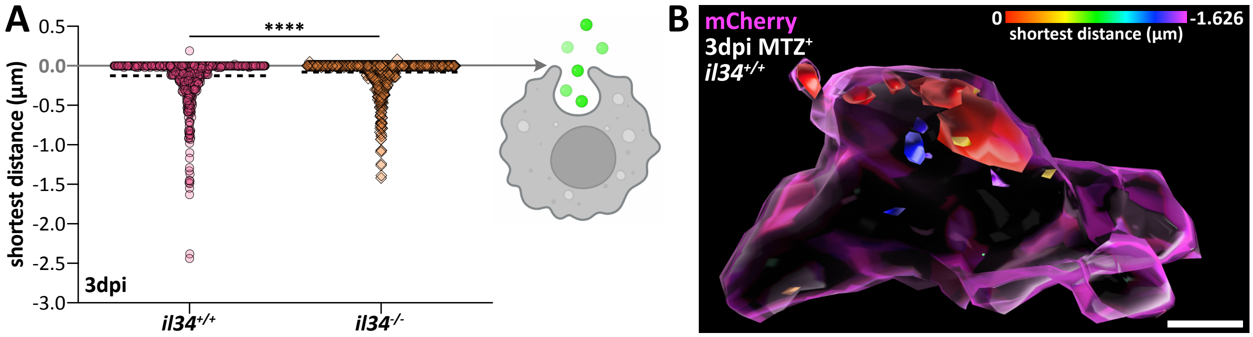
A
0.5
0.0
-0.5
-1.0
-1.5
-2.0
-2.5
-3.0
shortest distance (µm)
****
3dpi
il34+/+
il34-/-
B
mCherry
3dpi MTZ+
il34+/+
0
-1.626
shortest distance (µm)

**Figure S6**

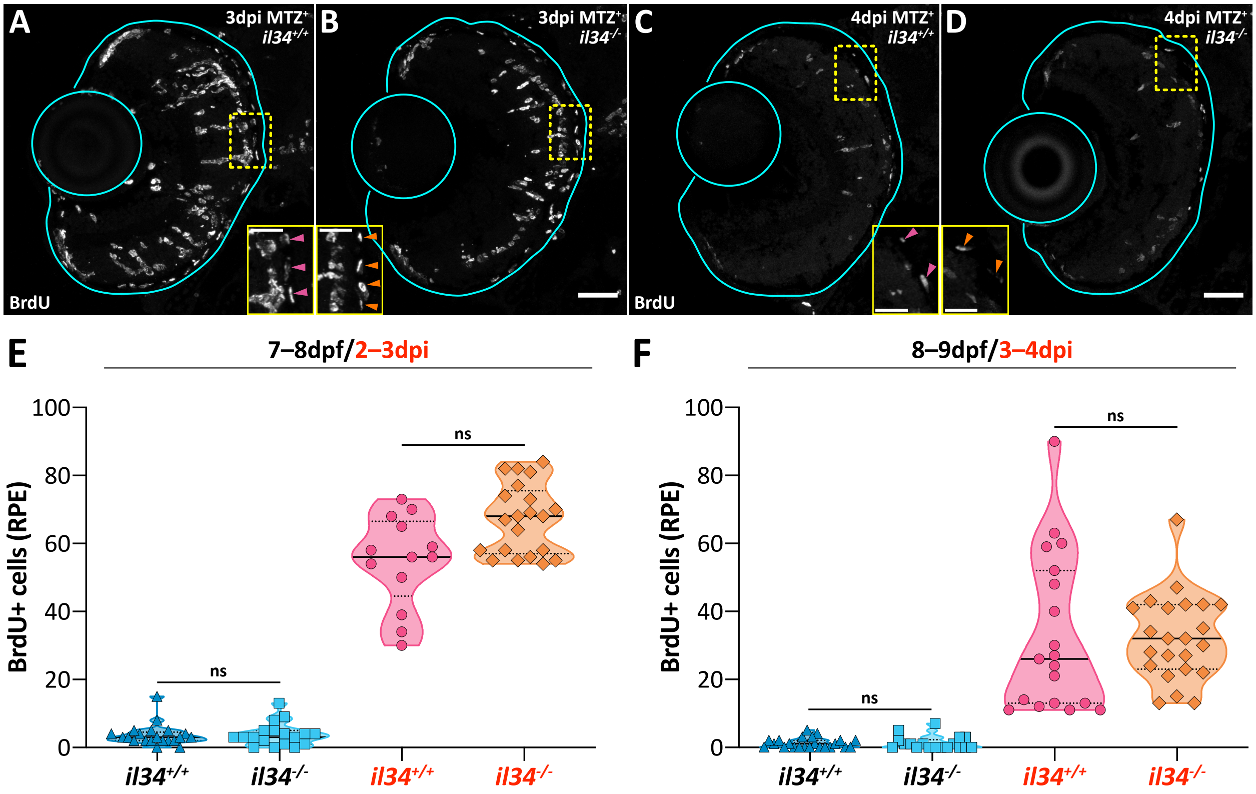
A
3dpi MTZ+
il34+/+
B
3dpi MTZ+
il34-/-
C
4dpi MTZ+
il34+/+
D
4dpi MTZ+
il34-/-
BrdU
BrdU
E
7–8dpf/2–3dpi
F
8–9dpf/3–4dpi
BrdU+ cells (RPE)
100
80
60
40
20
0
ns
il34+/+
il34-/-
il34+/+
il34-/-

**Figure S7**

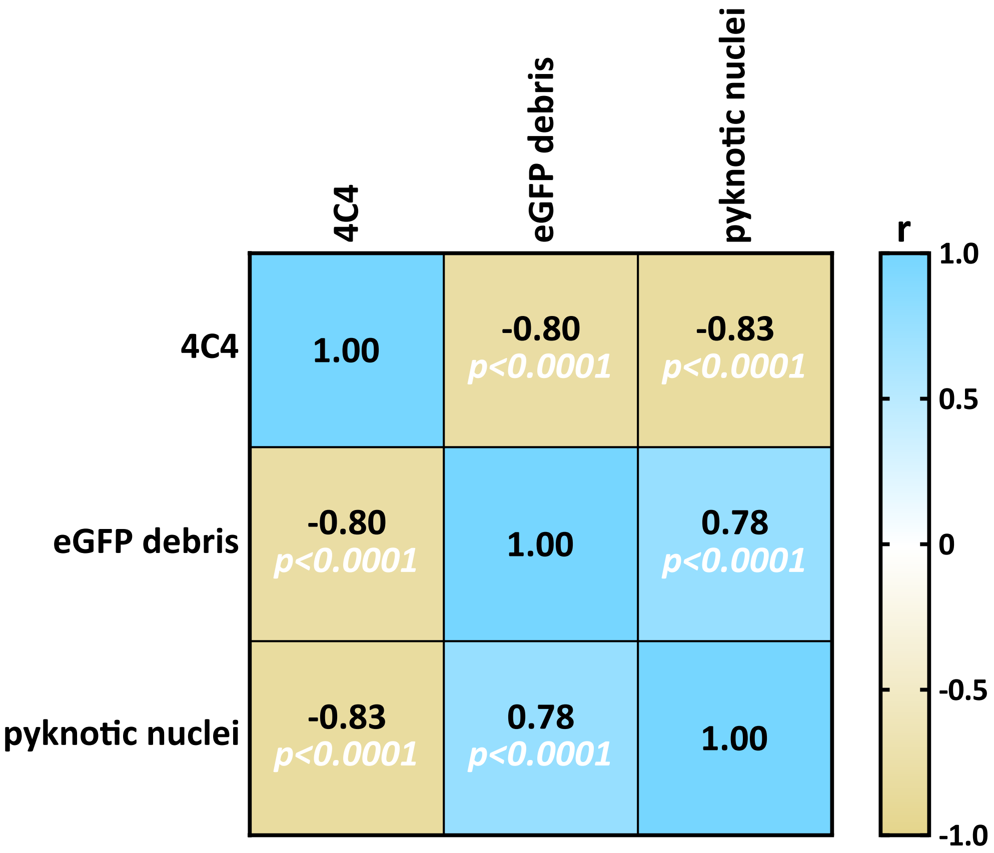
4C4
eGFP debris
pyknotic nuclei
r
1.0
0.5
0
-0.5
-1.0
4C4
eGFP debris
pyknotic nuclei
1.00
-0.80
p<0.0001
-0.83
p<0.0001
-0.80
p<0.0001
1.00
0.78
p<0.0001
-0.83
p<0.0001
0.78
p<0.0001
1.00
